# Antigen presentation dynamics shape the response to emergent variants like SARS-CoV-2 Omicron strain after multiple vaccinations with wild type strain

**DOI:** 10.1101/2022.08.24.505127

**Authors:** Leerang Yang, Matthew Van Beek, Zijun Wang, Frauke Muecksch, Marie Canis, Theodora Hatziioannou, Paul D. Bieniasz, Michel C. Nussenzweig, Arup K. Chakraborty

## Abstract

The Omicron variant of SARS-CoV-2 evades neutralization by most serum antibodies elicited by two doses of mRNA vaccines, but a third dose of the same vaccine increases anti-Omicron neutralizing antibodies. By combining computational modeling with data from vaccinated humans we reveal mechanisms underlying this observation. After the first dose, limited antigen availability in germinal centers results in a response dominated by B cells with high germline affinities for immunodominant epitopes that are significantly mutated in an Omicron-like variant. After the second dose, expansion of these memory cells and differentiation into plasma cells shape antibody responses that are thus ineffective for such variants. However, in secondary germinal centers, pre-existing higher affinity antibodies mediate enhanced antigen presentation and they can also partially mask dominant epitopes. These effects generate memory B cells that target subdominant epitopes that are less mutated in Omicron. The third dose expands these cells and boosts anti-variant neutralizing antibodies.

## Introduction

The emergence of viral mutants that escape from vaccine-imprinted immune memory is a major challenge for the development of vaccines against highly mutable viruses. In less than two years since effective vaccines became available, several SARS-CoV-2 variants of concern have emerged and spread. The Omicron variant harbors 32 mutations in the spike protein that enables it to escape from the majority of known monoclonal antibodies (Cao et al., 2022; Cele et al., 2022; Planas et al., 2022). Individuals vaccinated with two doses of mRNA vaccines encoding the spike protein of the original Wuhan strain have significantly lower neutralizing antibody titers against Omicron compared to the original strain. However, a booster shot (3^rd^ dose) of the same vaccine significantly increases protection against Omicron (Accorsi et al., 2022; Canaday et al., 2022; Lauring et al., 2022; Thompson et al., 2022).

After the booster, the peak neutralization titer increased roughly 3-fold against the wildtype (WT) Wuhan strain compared to the peak value after the 2^nd^ dose, but there was a 20-30 fold increase against Omicron (Garcia-Beltran et al., 2022; Muecksch et al., 2022; Schmidt et al., 2022). Thus, the booster shot increased the breadth of the resulting neutralizing antibodies in addition to restoring antibody titers that waned over time. The increase in breadth after the third dose has been attributed to the rise of antibodies targeting diverse epitopes in the receptor-binding domain (RBD), some of which are relatively conserved between the Wuhan and Omicron strains (Dejnirattisai et al., 2022; Muecksch et al., 2022).

SARS-CoV-2 RBD-binding neutralizing antibodies can be classified into different categories based on the regions/epitopes they target (Barnes et al., 2020a; Dejnirattisai et al., 2021). The most immunodominant epitopes lie in the ACE2 binding interface region (Garcia-Beltran et al., 2022; Greaney et al., 2021b, 2021a; Muecksch et al., 2022). Several human germline heavy chain genes exhibit high affinities for these epitopes (Yuan et al., 2020). Antibodies that develop from these germlines are highly enriched in the responses to both infection (Robbiani et al., 2020) and two doses of mRNA vaccination (Cho et al., 2021). The Omicron variant is highly mutated in the epitopes targeted by these antibodies, and therefore it can significantly evade the immune response generated after two doses of mRNA vaccines (Garcia-Beltran et al., 2022).

Some of the Omicron-neutralizing antibodies that develop after the third vaccine dose must target relatively conserved epitopes. These antibodies must be subdominant because they are not present in large titer after the second vaccine dose. Immunodominance during interclonal competition of germinal center (GC) B cells is not well-understood. It is thought to be shaped by a combination of factors that include the frequency and affinity of naïve B cells (Abbott et al., 2018; Amitai et al., 2020; Havenar-Daughton et al., 2018; Sangesland et al., 2019), antigen availability in the lymph node (Angeletti et al., 2019; Cirelli et al., 2019), re-activation of pre-existing memory B cells (Amitai et al., 2020; Wang et al., 2015) and epitope masking by pre-existing antibodies (Bergström et al., 2017; McNamara et al., 2020; Tas et al., 2022).

In this paper, we study the mechanisms that underlie how a third dose of the Wuhan strain’s spike changes the immunodominance hierarchy of the resulting antibody response. We first developed an *in-silico* model that integrates the expansion of memory B cells outside the GC (extra germinal centers or EGCs) with processes that occur in GCs, and also explicitly considers antigen presentation dynamics in lymph nodes after the first and subsequent shots of homologous vaccines. Our results show that antigen presentation in the GC differs markedly between the first and second shots, and this difference plays an important role in shaping the resulting memory B cell responses. Limited antigen availability in GCs after the first shot results in a memory response dominated by B cells that target immunodominant epitopes, which are heavily mutated in an Omicron-like strain. After the second shot, these memory B cells expand and differentiate into plasma cells outside GCs, dominating the antibody response. In secondary GCs, higher levels of antigen are available because antibodies generated after the first dose enable effective antigen presentation. The circulating polyclonal antibodies can also block some dominant epitopes more effectively than subdominant epitopes. These effects lead to an increase in memory B cells that target subdominant epitopes that are relatively conserved in an Omicron-like strain. After the third dose, these memory B cells expand and differentiate into plasma cells whose products confer better protection against emergent variants. These predictions from the *in-silico* model are consistent with our analyses of new and existing data obtained from individuals vaccinated with three shots of mRNA vaccines. Taken together, our results show that an interplay between antigen presentation dynamics and processes that occur in and outside GCs explain the change in antibody immunodominance hierarchy upon receiving the third shot of COVID mRNA vaccines. These insights shed new light on fundamental aspects of the nature of the recall response that are directly relevant to vaccine design. Our results also explain several recent observations linking different vaccine regimens to protection from Omicron (Regev-Yochay et al., 2022; Zhao et al., 2022).

### *In-silico* model for the humoral immune response

Our model incorporates four main aspects of the B cell and antibody responses: (i) antigen presentation on follicular dendritic cells (FDCs), (ii) activation of naive B cells and entry into GCs, (iii) affinity maturation in GCs and export of memory and plasma cells, and (iv) expansion of memory B cells and differentiation into plasma cells in EGCs (Fig. 1). A set of differential equations that extends a previous study (Tam et al., 2016) models antigen capture and transport. Stochastic agent-based models are used to simulate GC and EGC processes (Amitai et al., 2020; Van Beek et al., 2022; Wang et al., 2015). We consider four general classes of B cells: naive B cells, GC B cells, memory B cells, and plasma cells. At each incremental time step in the simulations, the probabilities of actions such as activation, selection, proliferation, mutation, differentiation, and apoptosis are calculated for the B cells, and these actions are then executed accordingly. 200 separate GCs are simultaneously simulated to mimic a secondary lymphoid organ (Jacob et al., 1991). The simulation is repeated 10 times for each vaccine dose. The average quantities thus calculated could be considered to be the typical population-level response. Descriptions of the computational model and the simulation algorithm are outlined below (see STAR Methods for further details of the model; Tables S1 and S2 for parameters used and Table S3 for detailed simulation algorithm).

**Figure 1.**
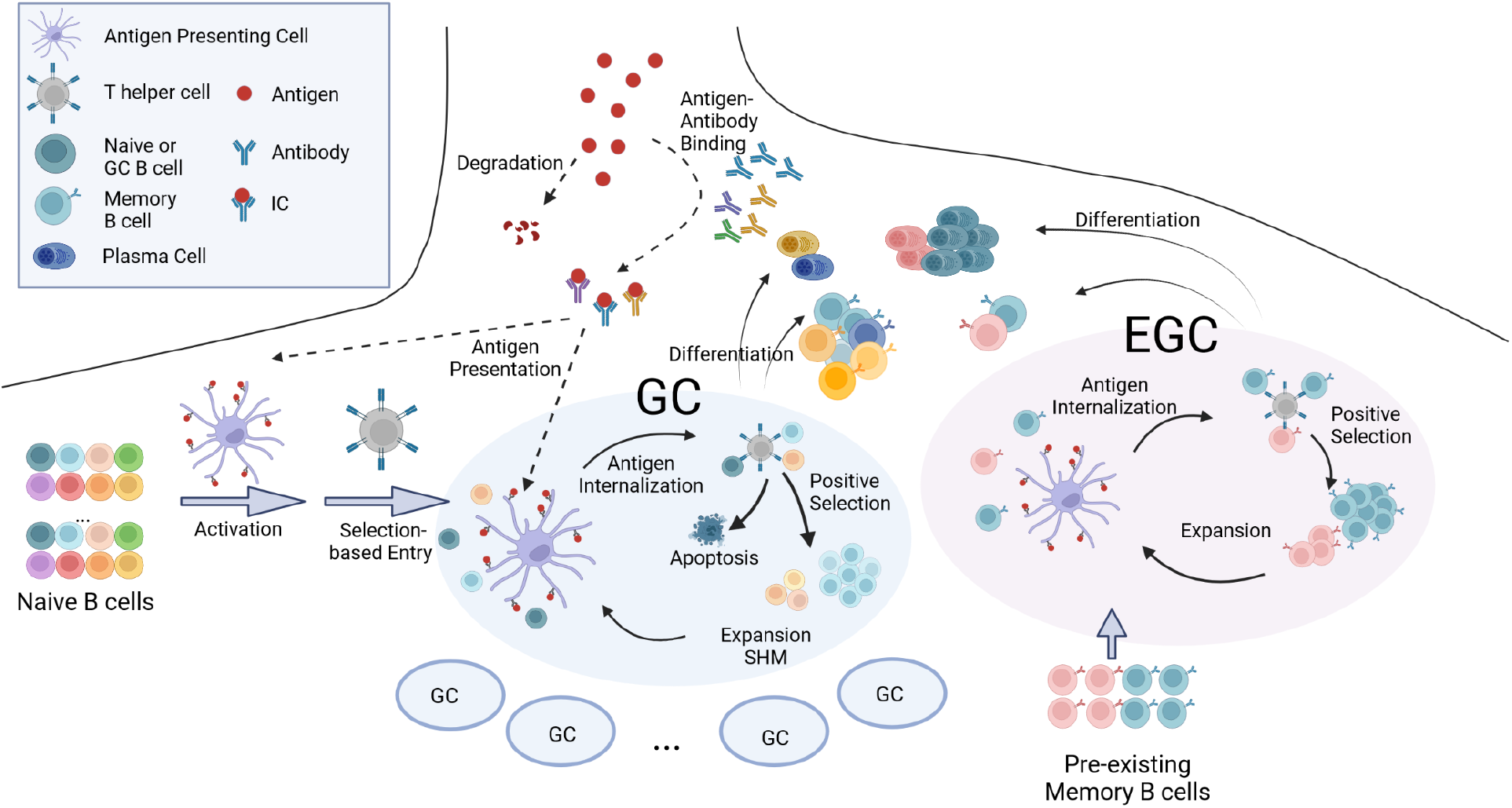
Schematic depiction of the In-Silico Model: The model integrates antigen presentation dynamics with processes in GCs and EGCs. Circulating antibodies help present antigen on FDCs. GC entry, GC B cell selection, replication and mutation, and differentiation of GC B cells into memory and plasma cells are considered. In the EGC, pre-existing memory cells undergo selection, proliferation, and differentiation without mutations. See also Figure S1. The figure was created with Biorender software.

#### Model for antigen presentation

Although mRNA vaccines induce antigen production *in-vivo*, the protein production rate decreases rapidly and exponentially (Pardi et al., 2015). So we model vaccination as injection of a bolus of antigen (Tam et al., 2016). Soluble and immune complex (IC) forms of the antigen rapidly reach dynamic equilibrium, with their relative amounts determined by the pre-existing antibody concentrations and equilibrium constants for antibody-antigen binding. The soluble antigen decays quickly but ICs are transported to FDCs where they decay with a much longer half-life. Upon immunization with a new antigen, small numbers of low-affinity circulating IgM antibodies are available to bind antigen. For subsequent immunizations, higher affinity antibodies generated by earlier GC/EGC processes are available to form ICs. The differential equations that describe IC formation and antigen presentation are coupled to the agent-based simulation of GC and EGC processes (parameters used, Table S1; simulation methods in STAR Methods).

#### Model for naive B cells and WT and variant strains

We model the distribution of germline-endowed affinities of naive B cells as an exponential distribution between *K*_*d*_ = 10^−6^*M* and 10^−8^*M*, where *K*_*d*_ is the dissociation constant. This is because a minimum affinity of about *K*_*d*_ = 10^−6^ *M* is required for activation (Batista and Neuberger, 1998), and rare, naive B cells with ~100-fold higher affinities can be found for antigens like SARS-CoV-2 (Feldman et al., 2021; Kuraoka et al., 2016). In our coarse-grained model, we group the few dominant epitopes on an antigen into a single “dominant” epitope and group the subdominant epitopes into a single “subdominant” epitope. The “dominant” epitope is targeted by a greater number of naive B cells and their affinities exhibit a longer high-affinity tail compared to the “subdominant” epitope (Fig. S1A; STAR Methods, Eqs. 2-5; parameters in Table S4).

Most immunodominant neutralizing epitopes on the SARS-CoV-2 spike protein are highly mutated in the Omicron variant (compared to WT) (Cao et al., 2022), while some subdominant epitopes are relatively conserved (Wang et al., 2022). Therefore, in our model, the dominant epitope is less conserved. A B cell has different affinities for the WT and the variant because the initial affinity and the effects of the mutations depend on the antigen (Fig. S1B, STAR Methods, Eqs. 6-7). The effect of mutations on binding affinities for the WT and the variant are drawn from correlated log-normal distributions so that ~5 % of affinity-affecting mutations are beneficial for each strain and most mutations are deleterious (Fig. S1C) (Kumar and Gromiha, 2006; Zhang and Shakhnovich, 2010). The level of correlation between the WT and variant distributions determines the fraction of mutations that will be beneficial for binding to both strains (Fig. S1D). We chose parameters so that about 72% and 19% of beneficial mutations increase affinities towards both strains for B cells that target subdominant and dominant epitopes, respectively. Our qualitative results are robust to changes in these parameters within reasonable ranges. Details of the simulation methods are in STAR Methods.

#### Model for germinal center entry of naive B cells

Naïve B cells continuously enter 200 GCs after activation and selection (Schwickert et al., 2007; Turner et al., 2017). At each time step, naive B cells internalize different amounts of antigen based on their binding affinity for the WT antigen and its availability (Batista and Neuberger, 1998; Fleire et al., 2006; Wang et al., 2015). Then, they stochastically get activated and compete for T helper cells for survival signals that allow GC entry (Lee et al., 2021; Okada et al., 2005; Schwickert et al., 2011). The probabilities for these entry events are determined by the amounts of internalized antigen (STAR Methods, Eqs. 10-13). The effect of memory B cell participation in GCs is studied by varying the fraction of pre-existing memory B cells added to the pool of naive B cells.

Selection stringency is an important factor in shaping B cell competition dynamics and thus the diversity of the response (Victora and Wilson, 2015). We studied the effects of changing the level of selection stringency and alternative models for antigen internalization to test the robustness of our qualitative results (STAR Methods, Eqs. 10 and 14).

#### Model for affinity maturation in germinal centers

For positive selection, GC B cells require activation by antigen capture (Luo et al., 2018; Shlomchik et al., 2019) followed by selection by T helper cells (Victora et al., 2010). In our model, GC B cells internalize antigen and are stochastically activated in the same way as the naive B cells. To model the competition for limited amount of T cell help, the birth rate of an activated GC B cell is determined by two factors: the amount of antigen it has captured relative to the average amount captured by all activated GC B cells, and the ratio between the number of T helper cells and activated B cells (STAR Methods, Eqs. 15-16). The number of T cells at a given time point is determined by a model that is fitted to a clinical observation in SARS-CoV-2 vaccinated subjects (STAR Methods, Eq. 17). All GC B cells also stochastically undergo apoptosis with a constant death rate (Mayer et al., 2017) (STAR Methods, Eq. 18).

With a probability, *p*_1_ each positively selected B cell exits the GC. It can differentiate into a plasma cell with probability *p*_2_, or become a memory cell. As discussed later, we studied varying *p*_1_ and *p*_2_. The remaining positively selected cells proliferate once. During a birth event, one of the two daughter cells mutates (Michael et al., 2002). A mutation leads to apoptosis (probability 0.3), no affinity change (probability 0.5), or a change in the mutation state of a randomly selected residue (probability 0.2) (Zhang and Shakhnovich, 2010). Details of the simulation methods are in STAR Methods.

#### Model for extra germinal center processes

Upon the second and third vaccination, an EGC response develops. EGCs select and expand pre-existing memory B cells without introducing mutations (Moran et al., 2019). The number of memory B cells peaked 1 week after the second dose in vaccinated subjects (Goel et al., 2021). Thus, although memory B cells may continue to be generated in EGCs, we terminate the EGC after 6 days in the simulation. The selection process is identical to that in the GCs, except that the number of T cells is assumed to be equal to the peak value to account for the fast kinetics of the EGC. Proliferating cells in the EGC differentiate into plasma cells with a high probability of 0.6 (Moran et al., 2018).

## Results

### Limited antigen availability after the first vaccine dose leads mostly to memory B cells that are descendants of naive B cells with high germline-endowed affinities for dominant epitopes

Our simulations show that after the first vaccine dose (Vax 1) only a small amount of antigen gets deposited and retained on FDCs (Fig. 2A). This is because soluble antigen decays rapidly and IgM antibodies with relatively low affinity for the new antigen form immune complexes. These results are consistent with images of antigen retention on FDCs in mouse and monkey lymph nodes after a first vaccine dose (Cirelli et al., 2019; Martin et al., 2021; Tam et al., 2016). In the first week after immunization, many naive B cells are activated and about 70 distinct cells enter each simulated GC (Fig. S2A), a result consistent with observations in mice (Tam et al., 2016; Tas et al., 2016).

**Figure 2.**
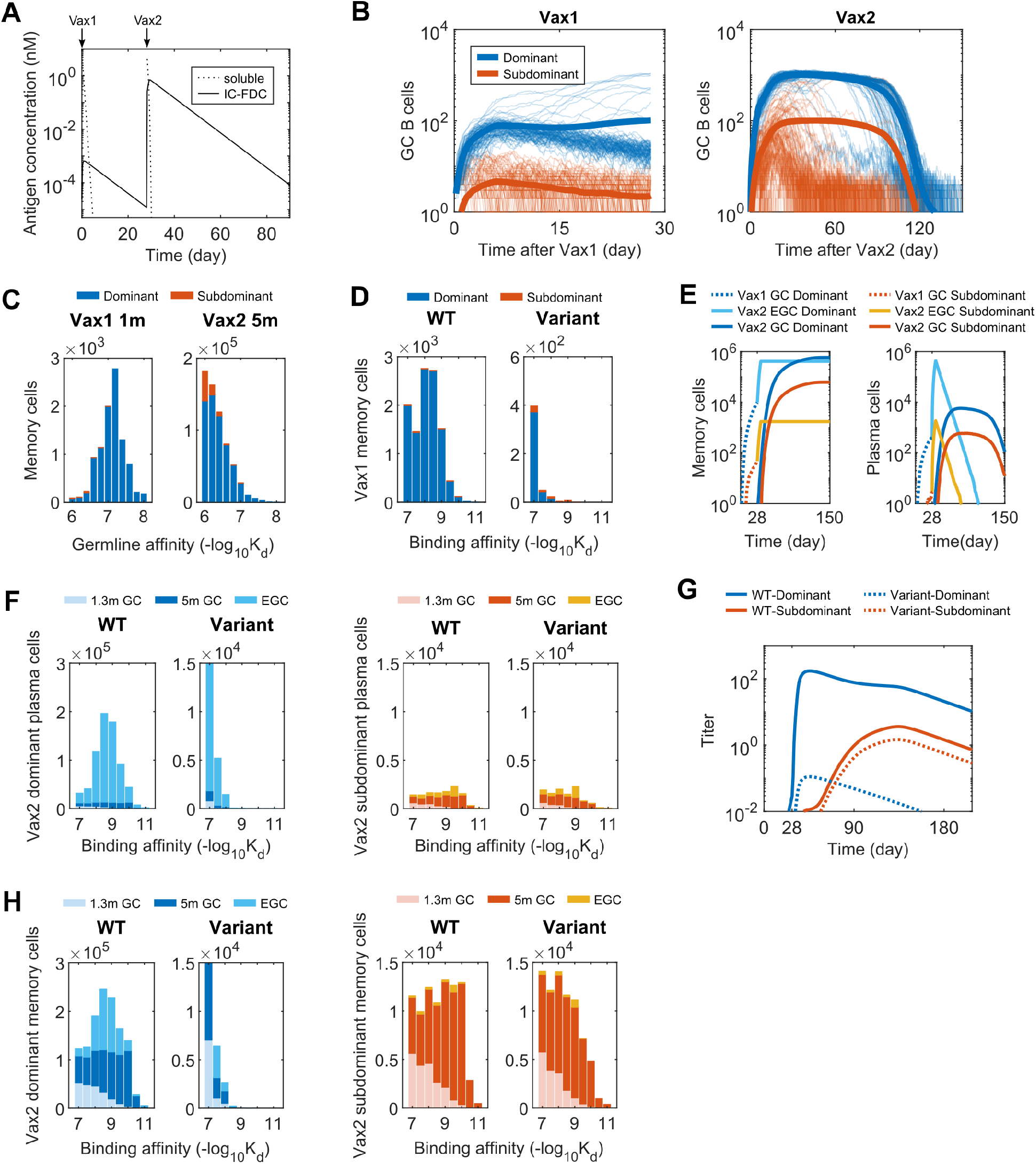
B cell and Antibody Responses after Vax 1 and Vax 2: **(A)** Concentrations of soluble antigen and immune complexes on FDCs. Vax 1 was administered on day 0 and Vax 2 on day 28. **(B)** Number of GC B cells that target dominant and subdominant epitopes after Vax 1 (Left panel) and Vax 2 (Right panel). 10 independent simulations of 200 GCs were performed for each case, and the bold curves show the mean values per GC. The other curves represent individual dynamic trajectories in 100 randomly selected GCs. **(C)** Histograms showing the distribution of WT-binding affinities of the germline B cell ancestors of GC-derived memory cells at 1m after Vax 1 (Left panel) and 5m after Vax 2 (Right panel). **(D)** Histograms showing the distribution of binding affinities of memory B cells against the WT and the variant at 1m after Vax 1. **(E)** Number of memory cells (Left panel) and plasma cells (Right panel) from GCs and EGCs after Vax 1 and Vax 2. Memory cells generated from Vax1 are expanded in the EGC and differentiate into plasma cells. New memory B cells and plasma cells are also generated from Vax 2 GCs. The plasma cells are short-lived and decay at a constant rate. **(F)** Histograms showing the distribution of binding affinities of plasma cells for the dominant (left panels) and subdominant (right panels) epitopes of the WT and the variant strains after Vax 2. GC-derived cells at 1.3m and 5m after vaccination and EGC-derived cells are shown. EGCs only last for six days, so no plasma cells are generated between 1.3m and 5m. Since plasma cells are short-lived, the data for a given time point shows all cells generated until that time. **(G)** Antibody titers after Vax 1 and Vax 2 that target the dominant and subdominant epitopes of the WT and the variant strains. Titers are calculated as the antibody concentrations divided by *K*_*d*_. **(H)** Histograms showing the distribution of binding affinities of memory cells for the dominant (left panels) and subdominant (right panels) epitopes of the WT and the variant strains after Vax 2. All histograms show distributions in terms of numbers of cells from 200 GCs, averaged over 10 simulations.

Given the low antigen availability, only high-affinity GC B cells can internalize enough antigen to have a high probability of receiving survival signals from T helper cells (Batista and Neuberger, 1998; Luo et al., 2018). Many B cells in the early GC fail to internalize enough antigen because their germline affinities are too low. GC B cells also develop deleterious mutations more frequently than beneficial ones (Kumar and Gromiha, 2006), which further reduces their chance of being positively selected. Thus, in many simulated GCs the B cell population begins to decline, which makes it even more unlikely that beneficial affinity-increasing mutations will evolve. In some GCs, beneficial mutations are not acquired soon enough to prevent GC collapse. In other GCs, B cells with high germline affinities proliferate and evolve beneficial mutations that increase antigen-binding affinities sufficiently to further proliferate, affinity mature and generate memory B cells. We find that ~75% of these memory B cells generated after Vax 1 originate from B cells with high germline affinities (−*logK*_*d*_ ≥ 7) even though they make up a small fraction (~0.06%) of naïve B cells, and these cells predominantly target dominant epitopes (Fig.s 2B and 2C). The genetic diversity in GCs is also limited (Fig. S2B) as a small number of high affinity B cells quickly dominate (Escarmís et al., 2006; Li and Roossinck, 2004). Thus, the memory response after Vax 1 is dominated by a small number of expanded clones (Fig. S2C), consistent with data from vaccinated humans (Cho et al., 2021). Since these B cells target immunodominant epitopes that are highly mutated in the variant, they exhibit limited cross-reactivity (Fig.s 2D and S2D).

Many observed neutralizing class 1/2 antibodies against WT SARS-CoV-2 that target dominant epitopes differ by only one or two mutations from the corresponding germline ancestors (Barnes et al., 2020b; Brouwer et al., 2020; Kreer et al., 2020; Yuan et al., 2020). Our results suggest that this is because the GC response after Vax 1 is dominated by a few expanded clones that originate from naive B cells characterized by relatively high germline affinity for the dominant epitopes. One or two mutations are sufficient for these B cells to successfully mature in GCs.

We chose a particular set of parameters (Table S3) to obtain the results shown in the main text, but we tested the robustness of this finding by varying the following key simulation parameters: the parameter that determines the relative importance of antigen availability for positive selection of GC B cells; parameters that characterize the naïve B cell repertoire and stringency of affinity-based selection. Our qualitative findings are robust across a wide range of these parameter values (Fig. S3A-D). Our results are also robust to using an alternative model for the selection of GC B cells (STAR Methods Eq. 14, Fig. S3E).

### Expansion and differentiation of existing memory B cells that target dominant epitopes control the antibody response after the second dose, while increased antigen availability in secondary GCs elicits memory B cells that target subdominant epitopes

After the second vaccine dose (Vax 2), the memory and plasma cell responses are determined by processes that occur in newly formed secondary GCs and in EGC compartments. Our choice of simulation parameters that characterize the relative numbers of plasma and memory cells that exit from the GCs and EGCs was informed by data from mice and humans (see the section on model). These data suggest that many short-lived plasma cells are rapidly produced in EGCs which then quickly decay, while GCs produce a relatively small number of plasma cells over longer times (Goel et al., 2021; Muecksch et al., 2022; Turner et al., 2021). The number of EGC and GC-derived memory B cells appear to be of similar orders of magnitude since the numbers of RBD-targeting memory cells are similar between ~1 month and ~5 months after Vax 2 (Goel et al., 2021; Muecksch et al., 2022). Our qualitative results are robust to parameter variations over wide ranges (Fig.s S3A-S3F).

Since EGCs select and expand the memory B cells generated in response to Vax 1 in an affinity-dependent manner (Fig. 2E), most of the plasma cells that differentiate from them target the dominant epitopes and have low cross-reactivity to the variant (Fig. 2F, S2D-E). Therefore, the WT antibody titer rapidly increases but not the variant titer (Fig. 2G). The number of plasma cells derived from secondary GCs is small compared to EGC-derived plasma cells (Fig. 2E-F) and has a limited contribution to the overall antibody titer after Vax 2, an observation consistent with original antigenic sin (Francis, 1960). That is, the antibody response to secondary immunization is dominated by the recall of previously generated responses.

After Vax 2, soluble antigen rapidly forms ICs with pre-existing high-affinity antibodies before it decays to low levels (Fig. 2A). Thus, we find a large difference in antigen availability after primary and secondary immunization, consistent with lymph node imaging of rhesus macaques (Martin et al., 2021). In the first week after immunization, a similar number of B cells join the GCs as in Vax 1 (Fig. S2A). The high amounts of antigen available on FDCs now allow lower affinity B cells that target subdominant epitopes to internalize antigen, proliferate, acquire beneficial mutations and compete with higher affinity cells for survival signals from helper T cells. Unlike Vax 1 GCs, this effect prevents secondary GC B cells from being completely dominated by high-affinity B cells that target dominant epitopes (Fig. 2B, S2B), and diverse memory B cells exit from the GCs (Fig. S2C). Since low-affinity naive B cells are much more common, they often ultimately outcompete the rare high-affinity naive B cells to take over GCs (Fig. 2C). Only ~7% of memory B cells descend from naive cells with high affinities after Vax 2 (−*logK*_*d*_ ≥ −7), in contrast to ~75% after Vax 1. By 5 months after Vax 2, large numbers of GC-derived memory B cells are produced, and they have higher affinities towards WT than the EGC-derived clones because of affinity maturation over time (Fig. 2H). Notably, by 5m after Vax 2, some subdominant epitope-targeting memory B cells also develop high affinities towards the variant (Fig. 2H, Fig. S2F).

We also studied the role of memory B cell re-entry into secondary GCs. We added different fractions of existing memory B cells to the naive B cell pool after Vax 2. We find that more memory B cell re-entry into GCs decreases the output of memory B cells that target subdominant epitopes (Fig. S4A). This is because most of the existing memory cells target dominant epitopes, and high-affinity memory B cells have a high chance of dominating the GC once they enter (Fig. S4B). These findings suggest that limiting memory B cell re-entry into the secondary GCs promotes the generation of memory B cells that target subdominant epitopes, and may be a mechanism that evolved to confer protection against future variants that may emerge (Mesin et al., 2019; Tas et al., 2022). Similar effects could result from alternative mechanisms such as the early export of predominantly low-affinity GC B cells as memory cells (Viant et al., 2020).

### Memory B cells generated in GCs after the second dose are expanded and differentiated in EGCs after the third vaccine dose to drive improved variant neutralization

After the third vaccine dose (Vax 3), existing memory B cells expand in the EGC and differentiate into plasma cells. A number of high-affinity memory B cells generated after Vax 2 target subdominant epitopes that are relatively conserved between the WT and variant strains (Fig. 2H). These cells differentiate into plasma cells with high affinity for the variant (Fig. 3A), Thus the antibody titer against the variant increases after Vax 3 (Fig. 3B). The fold-change in titer from 1.3 months post Vax 2 to 1 month post Vax 3 is greater for the variant than for the WT, consistent with serum responses in vaccinated humans (Muecksch et al., 2022; Schmidt et al., 2022). The breakdown of antibody titers based on epitope specificity shows that the variant-binding titer is driven by the subdominant epitope-targeting antibodies, while the WT-binding titer is still driven by the dominant epitope-targeting antibodies (Fig. 3B). The greater fold-change in variant-binding titer is therefore explained by the large increase in the number of subdominant memory B cells that emerge from Vax 2 GCs compared to that from Vax 1 GCs. Note that our results showing that neutralizing antibodies for the variant after Vax 3 are drawn from the existing memory pool after Vax 2 are consistent with clinical data showing that antibody sequences that neutralize Omicron after the third dose were present in the memory compartment after the second dose (Muecksch et al., 2022).

**Figure 3.**
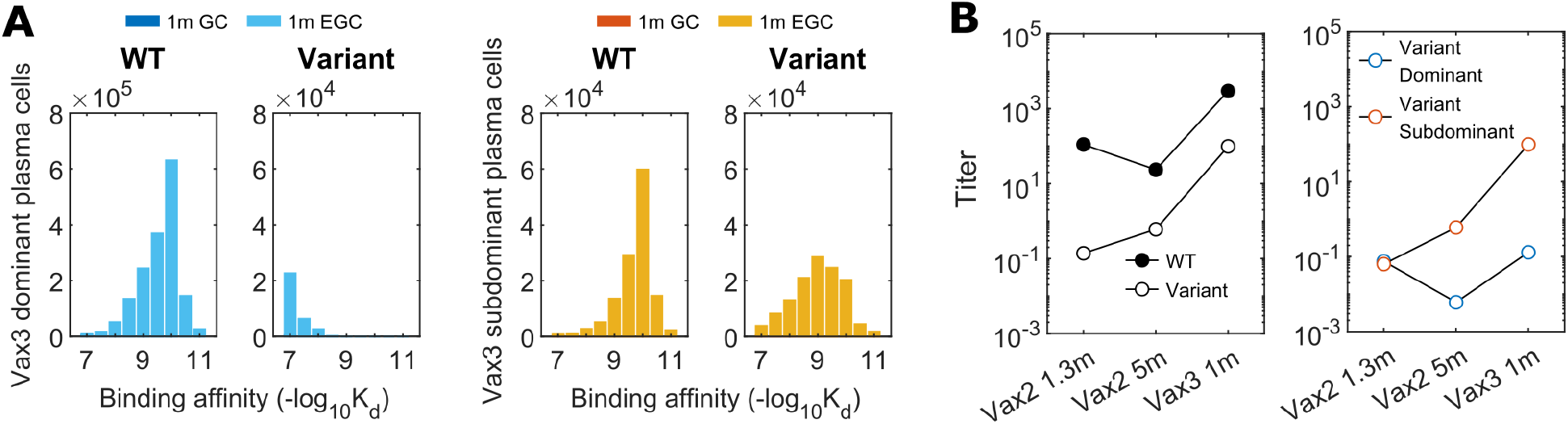
B cell and antibody responses after Vax 3: (A) Histograms showing the distribution of binding affinities of plasma cells targeting the dominant and subdominant epitopes of the WT and variant strains 1m after Vax3. At this point, almost all of the plasma cells are derived from the EGC. A substantial response to the subdominant epitope of the variant emerges. All histograms show distributions in terms of numbers of cells from 200 GCs, averaged over 10 simulations. (B) Comparison of antibody titers against the WT and the variant (left panel) and the epitope-specificity of the variant-targeting antibodies (right panel) at 1.3m after Vax 2, 5m after Vax 2, and 1m after Vax 3. The titer for antibodies targeting the subdominant epitope of the variant increases monotonically after 1.3m post Vax 2 because it has a very low value at early times. Titers are calculated as the antibody concentrations divided by *K*_*d*_.

### Analysis of sera from vaccinated humans is consistent with in-silico predictions

We explored the veracity of our *in-silico* predictions by analyzing data on sera obtained from individuals vaccinated with COVID-19 mRNA vaccines. Muecksh et al. sampled B cells from 5 uninfected individuals after the first, second, and third doses of the Moderna or Pfizer-BioNTech vaccines (Muecksch et al., 2022). The samples were collected an average of 2.5 weeks, 1.3 and 5 months, and 1 month after the first, second and third doses, respectively. We grouped sequences of 1370 B cells into clonal families and constructed a phylogenetic tree for each clonal family using Matlab’s seqlinkage function. If a phylogenetic tree contained two or more identical IGH sequences at the same time point or at different time points, we assumed that these clones were expanded in EGCs. The basis for this method is that EGCs expand memory cells with little to no mutations (Fig. S5A). This method is conservative as there is a low rate of mutation in EGCs (Moran et al., 2018). For this reason and because of under-sampling, we can identify only a small fraction of EGC-derived B cells. However, when tested against simulated data, we found the precision of our method for identifying EGC clones to be very high. From the simulation data in Figs. 2 and 3, we randomly sampled B cells from different time points as was done in experiments. We then applied the method described above, and found our identification method has a sensitivity of ~0.3 and a precision of ~0.9 for finding the EGC B cells (Fig. S5B). Bayesian analysis agrees with these estimates (STAR Methods, Fig. S5B). Sequences that were not EGC-derived were considered to be derived from GCs. Thus, we classified the sequences of B cells obtained after Vax 2 and Vax 3 as EGC-derived or GC-derived. The GC-derived cells were further classified as clones if clonally related sequences were observed and otherwise as singlets.

To test the *in-silico* results against clinical data, we determined the neutralization activity of antibodies derived from the sequences classified as EGC-derived and GC-derived. We combined existing data (Muecksch et al., 2022) with new measurements of neutralization activities for some of the sequences that our analyses identified as EGC-derived. The new measurements were carried out using the methods described before (Muecksch et al., 2022). The neutralization activities (IC_50_) of 112 antibodies derived from B cells were measured against the Omicron RBD. Nine EGC-derived B cells were identified from samples collected after Vax 2. Other B cells sampled 5 months after Vax 2 were labeled as GC-derived clonal families or singlets. The EGC-derived clones have a much higher IC_50_ than the likely GC-derived clones or singlets in terms of the mean and the maximum (Fig. 4A), indicating their low potency. The geometric mean of the GC-derived clones and singlets is 341 ng/mL which is much lower than the 919 ng/ml for EGC clones (p=0.00027). This result agrees with the *in-silico* prediction that GC-derived B cells exhibit better omicron neutralization titers than the EGC-derived B cells after Vax 2 (Fig. 2H, Fig. S2E,F). We note that five of the nine EGC-derived B cells after Vax 2 also did not neutralize the WT (Table S4).

**Figure 4.**
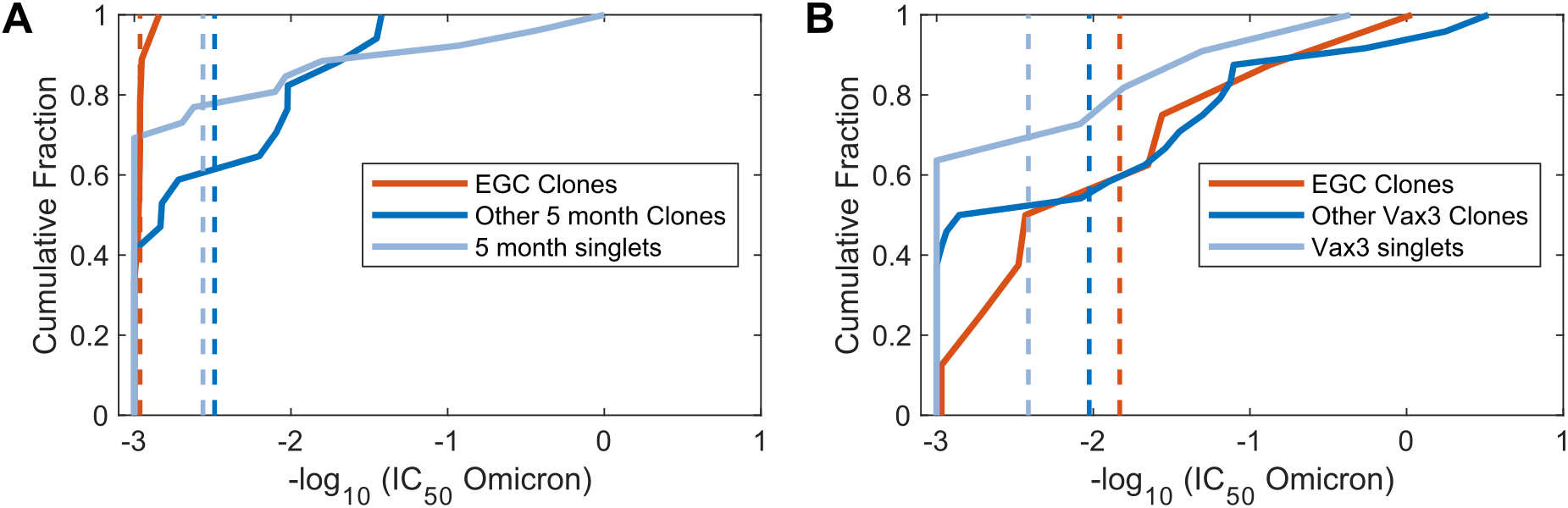
Omicron neutralization potency of monoclonal antibodies that are inferred to originate in EGCs and GCs, derived from vaccinated humans: (A) The cumulative distributions of omicron neutralization titers (IC_50_) of B cells and their antibodies sampled after Vax 2. Based on the sequence analysis (see text), the B cells have been classified as those identified to be derived from EGCs (red curves), other clonal families (blue curves), or singlets (light blue curves). Dashed lines indicate mean values. Because the EGCs are short-lived and the distributions were identical, EGC B cells collected 5 months after Vax 2 were combined with EGC B cells collected 1.3 months after Vax 2. **(B)** Similar data as in panel A for cells sampled 1m after Vax 3. A statistical comparison of the distributions shown in Panels A and B is noted in the text.

8 EGC-derived B cells were identified after Vax 3. Fig. 4B shows that the IC_50_ of EGC clones improved from a geometric mean of 919 ng/mL after Vax 2 to 68 ng/mL after Vax 3 (p=0.0035, STAR Methods). Comparing Figs. 4A and 4B shows that the geometric mean of IC_50_ values for EGC-derived antibodies after Vax 3 is more similar to the GC-derived ones after Vax 2 (341ng/ml) than the EGC-derived clones after Vax 2 (919 ng/ml). This is consistent with our *in-silico* predictions (Fig. 3A and Fig. 2H), which show that the EGCs formed after Vax 3 expand the subdominant and cross-reactive memory B cells generated after Vax 2.

### Epitope masking by polyclonal antibodies amplifies the increase in subdominant responses, but increased antigen availability plays a key role

Circulating antibodies can mask their corresponding epitopes, allowing GC B cells that target other epitopes to evolve. It has been speculated that masking of dominant epitopes by circulating antibodies may drive the diversity increase of memory B cells upon repeated mRNA vaccinations (Cameroni et al., 2022; Kotaki et al., 2022; Muecksch et al., 2022). Given the reported serum RBD-targeting antibody concentrations and affinities after mRNA vaccination (Demonbreun et al., 2021; Macdonald et al., 2022), the extent to which antibodies mask their corresponding epitopes can be calculated assuming dynamic equilibrium (Zhang et al., 2013). Such a calculation suggests that epitope masking will not be important after Vax 1 because of low antibody titer, but 2 weeks after Vax 2, antibodies will mask ~99% of the epitopes (Fig. S6A). If the dominant and subdominant epitopes do not overlap, then epitope masking selectively lowers the effective dominant epitope concentrations by ~100-fold. In our simulations, this causes subdominant B cells to monopolize the secondary GC response (Fig. S6B-C), consistent with experimental studies using monoclonal antibodies to block immunodominant epitopes (Bergström et al., 2017; McNamara et al., 2020; Xu et al., 2018).

However, antibodies developed after mRNA vaccination are highly polyclonal and target many overlapping epitopes. Class 1 and 2 neutralizing antibodies that dominate early antibody responses bind to the ACE2 binding motif (Barnes et al., 2020a; Cao et al., 2022). Rare Class 3 and 4 neutralizing antibodies target relatively conserved peripheries of the RBD (Cao et al., 2022; Muecksch et al., 2022). Some antibodies span multiple classes. Reanalysis of data from Muecksch et al. (Muecksch et al., 2022) shows class 1, 2, 3, and 1/4 antibodies interfere with 20-50% of the polyclonal antibodies across all time points (Fig. 5A). These data suggest that both dominant and subdominant epitopes will likely be partially blocked by serum polyclonal antibodies due to overlap between epitopes.

**Figure 5.**
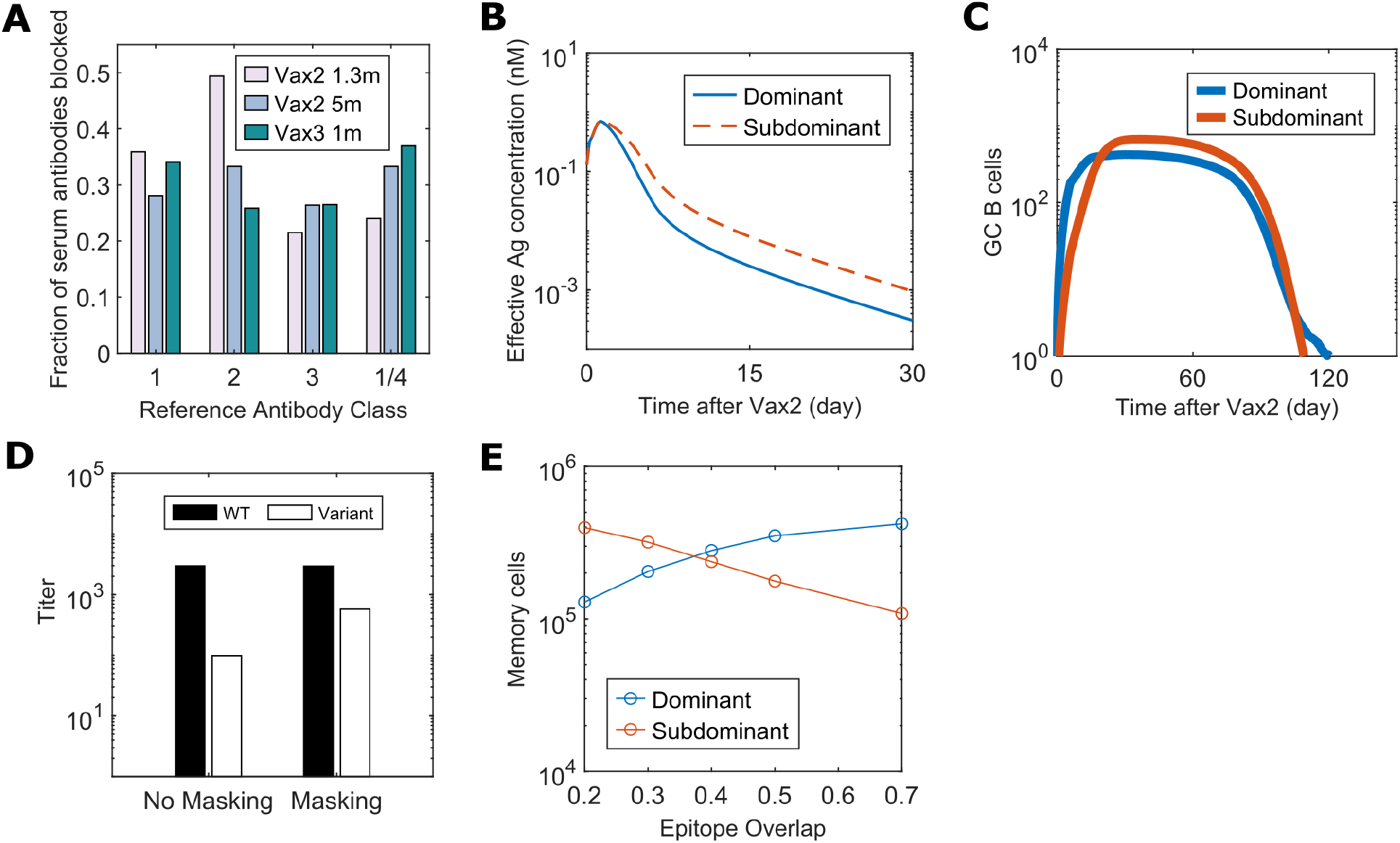
Role of epitope masking on immunodominance hierarchy: (**A)** Fraction of antibodies derived from human serum responses that blocked the binding of four reference antibodies (class 1, 2, 3, and 1/4) that target different regions of the SARS-CoV-2 RBD. Data from Muecksch et al. were reanalyzed (Muecksch et al., 2022). **(B)** Epitope-dependent effective antigen concentrations when there is epitope masking with 30% of epitope overlapping. **(C)** Number of GC B cells that target dominant and subdominant epitopes after Vax 2 with 30% epitope overlap. **(D)** Comparison of antibody titers at 1m after Vax 3 between simulations with no epitope masking (“No Masking”) and epitope masking with 30% of epitope overlap (“Masking”). Titers are calculated as the antibody concentrations divided by *K*_*d*_. **(E)** Number of dominant and subdominant memory B cells at 5m Vax 2 when the degree of epitope overlap is varied in simulations.

Therefore, we studied an epitope masking model where a fraction of antibodies targeting dominant epitopes can also block subdominant epitopes, and *vice versa*. When this fraction (epitope overlap) is 30%, the antigen availability advantage for subdominant B cells is relatively small (Fig. 5B). But this moderate effect amplifies the subdominant B cell response from the secondary GCs (Fig. 5C). Compared to the case without epitope masking, the antibody titer for the variant further increases after Vax 3, without much difference in the WT titer (Fig. 5D). Thus, our model suggests that epitope masking from polyclonal responses can enhance subdominant B cell responses. Epitope masking likely plays a significant role in the increase in class 3 and 4 neutralizing antibodies (that bind to the RBD periphery) after Vax 3 (Muecksch et al., 2022).

Well-conserved, subdominant epitopes also exist on the ACE2 binding motif that is targeted by class 1 and 2 antibodies (Wang et al., 2022). These epitopes overlap significantly with the epitopes targeted by dominant class 1 and 2 antibodies because antibody footprints typically cover most of the ACE-2 binding motif (Barnes et al., 2020a; Lan et al., 2020). Our calculations show that the effect of epitope masking on immunodominance decreases with an increase in the degree of overlap between dominant and subdominant epitopes (Fig. 5E). Analysis of 43 Omicron-neutralizing antibodies isolated from humans after Vax 3 showed that 63% of them were class 1/2 antibodies (Andreano et al., 2022). Class 1/2 antibodies derived from immunodominant germlines were prevalent in the early Vax 2 response, but the Omicron-neutralizing class 1/2 antibodies were derived mostly from subdominant germlines that were rarely observed 1.3m after Vax 2. These subdominant antibodies were significantly mutated 5m after Vax2, suggesting their development in secondary GCs (Andreano et al., 2022). Since these Omicron-neutralizing antibodies target epitopes that likely overlap significantly with the epitopes of the initially observed class 1/2 antibodies derived from dominant germlines, epitope masking alone cannot explain their rise. Increased antigen availability after Vax 2 likely plays a key role in promoting their emergence.

## Discussion

We studied the effects of repeated immunization with a WT vaccine on antibody responses to a highly mutated variant, such as the Omicron strain of SARS-CoV-2. Our findings shed new light on fundamental aspects of the humoral immune response, and can guide the design of vaccination strategies that aim to elicit broadly protective responses against mutable viruses.

After Vax 1, the amount of antigen presented on FDCs is limited (Fig. 2A). This is because soluble antigen decays quickly and only weakly binding IgM is available to form ICs. The limited antigen availability during GC reactions strongly promotes the evolution of memory B cells and antibodies that bind to immunodominant epitopes (Fig. 2B). These B cells are largely derived from naive B cells that bind to these epitopes with high germline affinity or can acquire high affinity via a small number of mutations (Fig. 2D). Upon receiving Vax 2, memory B cells generated by GCs after Vax 1 are rapidly expanded and they differentiate into plasma cells that secrete antibodies (Fig. 2E-F). Thus, the antibodies produced after Vax 2 largely target immunodominant epitopes. These epitopes are highly mutated in Omicron, and so Omicron neutralizing titers are low (Fig. 2G). These *in-silico* results are consistent with data showing the dominant antibodies produced after the first two doses have few mutations (Muecksch et al., 2022).

After Vax 2, higher amounts of antigen are displayed on FDCs because higher affinity antibodies produced after Vax 1 can form ICs efficiently (Fig. 2A). The higher antigen availability allows memory B cells that target subdominant epitopes to emerge despite their weaker germline affinities (Fig. 2B-C). These epitopes are relatively conserved between the WT and Omicron strains. After Vax 3, these memory B cells are expanded in EGCs resulting in increased Omicron neutralizing antibody titers (Fig. 3B). This is consistent with data showing that the Omicron-neutralizing antibodies present after Vax 3 existed in the memory pool after Vax 2. Importantly, our *in-silico* predictions are consistent with sequence and neutralization data that we obtained from vaccinated individuals and analyzed (Fig. 4).

Our results also provide mechanistic insights into the effects of the timing of booster shots on the ability to develop Omicron-neutralizing antibodies. A group of subunit vaccine ZF2001 recipients who received Vax 3 only 1 month after Vax 2 were less likely to develop Omicron neutralizing antibodies than the group with a 4 month interval (56% vs. 100%) (Zhao et al., 2022). Our model predicts (Fig. 6) that when Vax 3 is given 1.3 month after the second dose (“Vax3-Short”), the subdominant epitope-targeting antibody titer is low. Most of the memory cells that have high affinities 1.3 month after Vax 2 are EGC-derived and thus target the dominant epitope (Fig. 2H). Also, even subdominant GC-derived memory B cells have a relatively low affinity towards the variant due to limited time for affinity maturation (Fig. 2G). As a result, receiving Vax 3 1.3 month after Vax 2 will mostly expand B cells with low cross-reactivity. But 4 months after Vax 2, more affinity maturation allows B cells with higher affinity for subdominant epitopes to develop, which is consistent with the observation that the number of mutations increases significantly between 1.3 months and 5 months after Vax 2 (Muecksch et al., 2022). The memory B cells available 4 months after Vax 2 can be expanded in EGCs after Vax 3 to result in better Omicron neutralizing capability.

**Figure 6.**
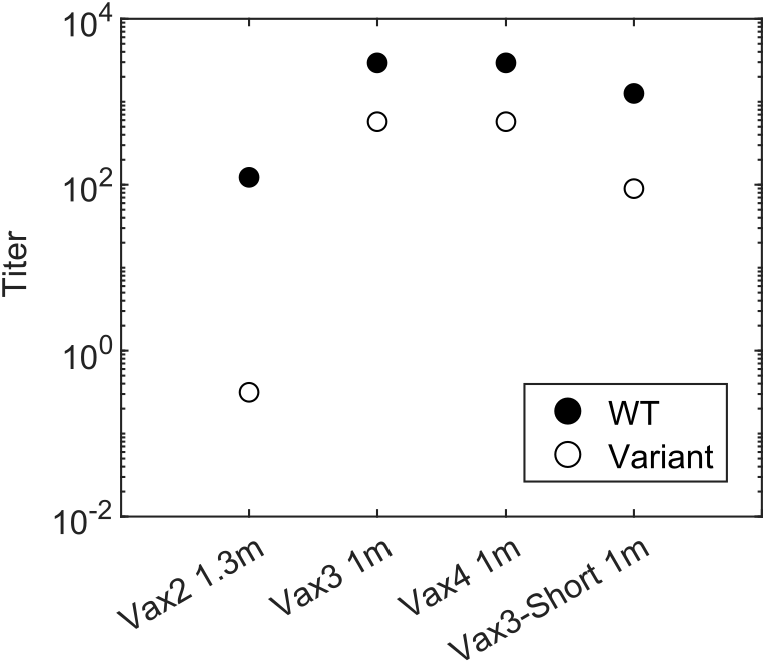
Comparison of antibody titers for different vaccination regimens: Antibody titer elicited by different vaccination regimens. “Vax4” refers to the case when a second booster dose was given 5m after Vax3. “Vax3-Short” refers to the case when Vax 3 was given 1.3m after Vax 2 instead of the standard 5m interval. To study how epitope masking may affect the second booster (Vax 4), all cases were simulated with epitope masking and 30 % epitope overlap. Titers are calculated as the antibody concentrations divided by *K*_*d*_.

Our results show that epitope masking can amplify subdominant B cell responses after booster shots even when a polyclonal response is elicited after Vax 1. This effect is especially important when the dominant and subdominant epitopes do not overlap much. Epitope masking likely plays an important role in the increase in class 3 and 4 antibody titers after Vax 3 because their epitopes do not overlap as much with dominant class 1 and 2 antibodies (Muecksch et al., 2022). However, when the extent of overlap between dominant and subdominant epitopes is higher, the importance of epitope masking diminishes (Fig. 5E). Dominant and subdominant epitopes targeted by class 1 and 2 antibodies have significant overlap. The observation of many Omicron-neutralizing subdominant class 1 and 2 antibodies after Vax 3 (Andreano et al., 2022) points to the importance of high antigen availability in promoting the emergence of subdominant responses upon boosting.

Regev-Yochay et al. reported that a fourth dose of an mRNA vaccine restored the antibody titer against Omicron to a level similar to the peak response after Vax 3, but unlike Vax 3 it did not further boost the titer compared to the previous dose (Regev-Yochay et al., 2022). Results from our model are consistent with this finding (Fig. 6). The mechanistic explanation is that GCs formed after Vax 3 do not benefit further from increased antigen availability compared to the GCs formed after Vax 2. Moreover, antibodies that target subdominant epitopes are available in higher titers soon after Vax 3 and they can mask these epitopes. Therefore, masking immunodominant epitopes confers less of an advantage to the subdominant B cells in GCs formed after Vax 3 compared to those formed after Vax 2. Thus, similar or fewer subdominant GC B cells develop after Vax 3. However, overall antibody titer after the fourth dose is still similar to Vax 3 because both GC and EGC-derived memory cells generated after Vax 3 are expanded.

Our results may also have implications for efforts to elicit broadly neutralizing antibodies (bnAbs) against HIV by sequential immunization with variant antigens (Escolano et al., 2016, 2021; Wang et al., 2015). This approach aims to focus the B cell response on a conserved target epitope. Higher antigen availability and masking of the conserved epitope after booster shots will likely promote the evolution of off-target responses in secondary GCs, consistent with observations in macaques (Escolano et al., 2021). These effects may be especially significant when the conserved target epitope is quite distinct from the diverse variable regions, as is the case for some epitopes targeted by bnAbs against HIV and the conserved epitope in the stem of influenza’s spike (Klein et al., 2013; Wu and Wilson, 2020).

Although memory B cells participating in secondary GCs can help protect against closely-related variants, our results show that these memory B cells can limit epitope diversification and adversely impact the ability to protect against variants that differ more significantly from the WT strain. This is because the affinity advantage of memory cells can allow them to dominate GCs. We note also that higher antigen availability and epitope masking may underlie recent observations in mice showing that memory B cells are not highly represented in secondary GCs (Mesin et al., 2019).

We hope that our results and mechanistic insights will motivate other fundamental studies into how the humoral immune response is influenced by antigen presentation dynamics. For example, it will be interesting to explore whether strategies to modulate antigen availability such as slow antigen delivery and immunization with immune complexes or particulate immunogens may help mitigate unwanted immunodominance hierarchies (Cirelli et al., 2019; Moyer et al., 2020; Pauthner et al., 2017).

## Acknowledgments

This research was supported by NIH grant # U19AI057229 and by the Ragon Institute of MGH, MIT, and Harvard (LY, MVB, AKC). MCN was supported by NIH grant # AI037526-27. Z.W was supported in part by grant # UL1 TR001866 from the National Center for Advancing Translational Sciences (NCATS, National Institutes of Health (NIH) Clinical and Translational Science Award (CTSA) program. PDB and MCN are Howard Hughes Medical Institute Investigators.

## Supplemental Figures and Tables

**Figure S1.**
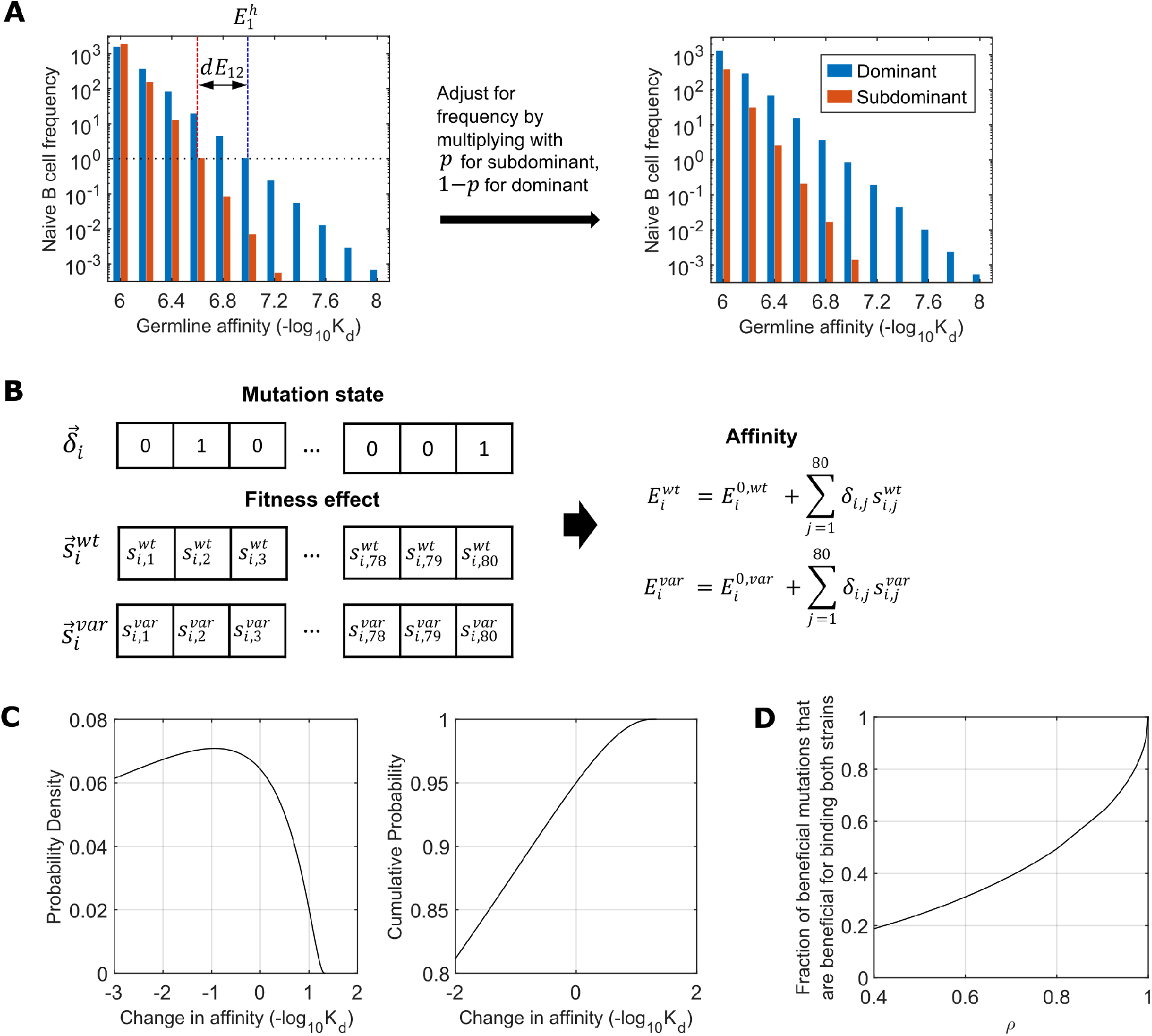
Simulation Details for B cells and 2-epitope model: **(A)** Distribution of germline-endowed affinities of naïve B cells, parameterized by 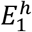, *dE*_12_, and *p*. Left panel) 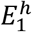 and *dE*_12_ specifies the slopes of the distributions. The naïve B cells can have germline affinities between 6 and 8 at intervals of 0.2. 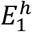 is the affinity at which the frequency of naïve B cells that target the dominant epitope would be 1 per GC, assuming there are 2000 B cells distributed according to a geometric distribution. It thus specifies the slope of the distribution for the B cells that target the dominant epitope. Analogously, 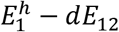 specifies the slope for the B cells that target the subdominant epitope. Right panel) The distributions are adjusted based on the parameter *p*. The fraction of all naïve B cells that target the dominant and subdominant epitopes are 1 − *p* and *p*, respectively. The distributions from the left panel are multiplied by these values to obtain the actual naïve B cell frequencies. **(B)** Schematics showing how the binding affinities against the WT and the variant strains are determined for a given B cell, *i*. The mutation state vector, 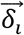, is initially a string of zeros, and some residues are mutated to ones during affinity maturation. The effects of a mutation of each residue on the WT and variant affinities (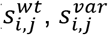 for residue, *j*) are drawn from a correlated probability distribution. The binding affinities 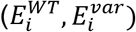 are sums of the initial affinities 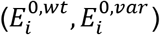 and the effects of mutated residues. **(C)** Marginal probability density function and cumulative distribution function for both the random variables 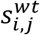 and 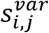. That is, they show probabilities of how a mutation of one residue from 0 to 1 will change the binding affinities. Although a single mutation will contribute differently to the WT and variant affinities, statistically for both variants ~5% of all mutations increase affinity, and the best beneficial mutations increase the affinity by ~10 fold. **(D)** The fraction of mutations that increase the affinity against the WT, which also increase the affinity against the variant. 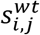 and 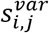 are drawn from correlated distributions parameterized by *ρ*, so that as *ρ* increases, mutations that are beneficial for binding both strains become more common. Thus, *ρ* represents the level of conservation of the epitope between the WT and the variant.

**Figure S2.**
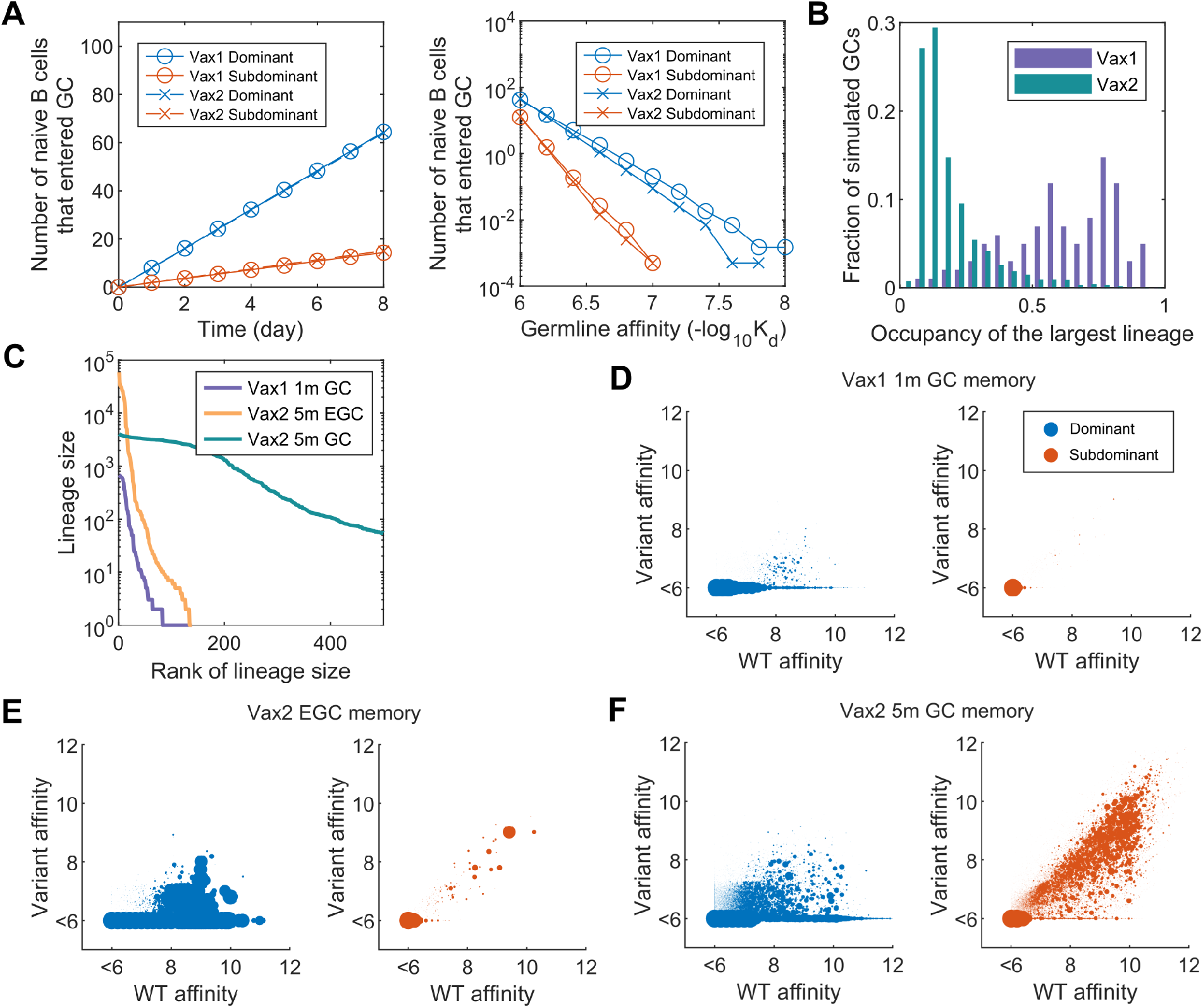
Details of Vax1 and Vax2 B cell response: **(A)** Left panel) Mean number of naïve B cells that enter a GC over time. The model allows similar numbers of naive B cells to enter GC after Vax 1 and Vax 2. Right panel) Germline affinities of B cells that have entered GC by day 8. High-affinity naïve B cells are more favored to enter GC after Vax 1 than after Vax 2 because of low antigen availability. However, since only a small number of such cells exist, most naïve B cells that enter GC are low-affinity B cells in both cases. Thus, the profiles of naïve B cells that enter GC after Vax 1 and Vax 2 are similar, as shown in the left panel. This model is conservative because higher antigen availability after Vax 2 could increase the number of naïve B cells that enter GC, which would strengthen the finding of greater B cell diversity from the Vax 2 response. **(B)** Histogram showing the distribution of the fraction of GC B cells that belong to the single largest lineage at 14 days after Vax 1 and Vax 2. Most of the Vax 1 GCs are already dominated by a single lineage at this time, in contrast to the Vax 2 GCs. **(C)** Number of memory cells from the same lineages, shown in the order of largest to smallest lineages. A few largest lineages dominate the Vax 2 EGC response, like Vax 1 GC response. In contrast, diverse lineages of similar sizes make up the Vax 2 GC response. The result shown is from a single simulation of 200 GCs. **(D-F)** Cross-reactivity of memory B cells derived from GCs and EGCs. The areas of the markers scale with the number of cells that have identical affinities. (D) GC at 1m after Vax 1, (E) EGC after Vax 2, (F) GC at 5m after Vax 2. Only a very small number of subdominant memory cells are generated after Vax 1, and they undergo limited expansion in EGC after Vax 2. In contrast, diverse subdominant B cells are produced in Vax 2 GCs, some of which have high affinities towards the variant.

**Figure S3.**
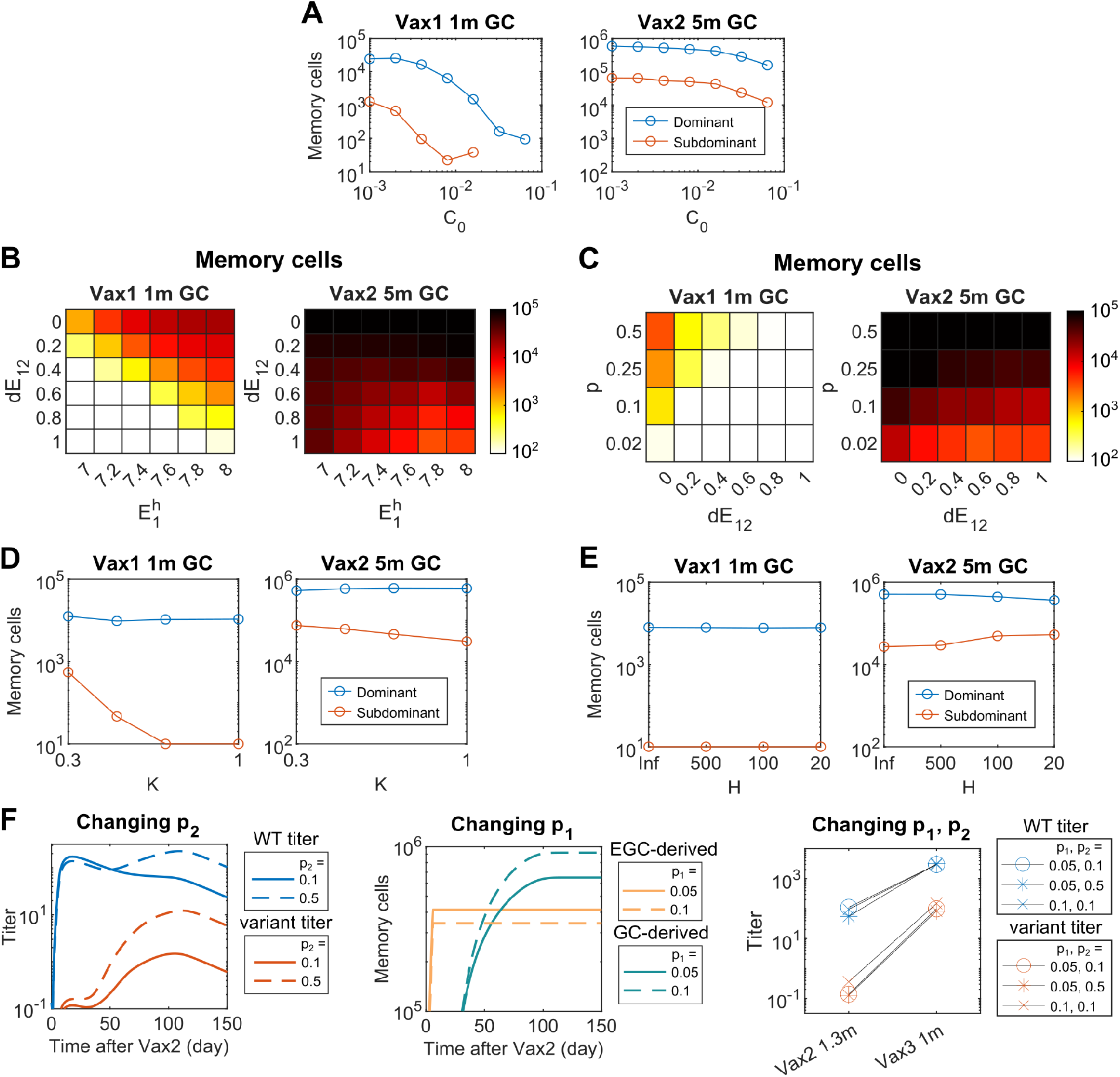
Parameter sensitivity analysis: Number of memory B cells derived from GCs at 1 month after Vax 1 and 5 months after Vax 2 that target dominant and subdominant epitopes, when various simulations parameters are changed. **(A)** The reference antigen concentration, *C*_0_, is varied. Decreasing *C*_0_ makes B cells easily activated even when the antigen concentration is small. The quantitative difference between the number of subdominant memory cells after Vax 2 and Vax 1 is largest when *C*_0_ is large; that is, when the importance of antigen concentration is high. However, the qualitative trend that more subdominant B cells are generated after Vax 2 is robust across ~2 orders of magnitude variation in *C*_0_. **(B-C)** Parameters that characterize the affinity distribution of naïve B cells are varied. 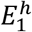 and *dE*_12_ are varied in (B), *p* and *dE*_12_ are varied in (C). For some parameter values, especially small *dE*_12_ and large *p*, some subdominant memory cells develop after Vax 1. However, for all parameter values, the number of subdominant B cells greatly increases after Vax 2, showing the robustness of our findings. **(D)** Parameter *K*, which controls the stringency of selection, is varied. Both after Vax 1 and Vax 2, more subdominant B cells develop when selection is permissive (small value of *K*). For all values of *K* tested, the number of subdominant B cells greatly increases after Vax 2 compared to Vax 1. **(E)** An alternative model of antigen capture is used, and the parameter *H* is varied. With this model, the amount of antigen captured by B cells saturates if the product of B cell affinity and antigen concentration is much greater than *H*. The original model is equivalent to infinite *H*. The qualitative finding is robust to changes in the model. Quantitatively, slightly more subdominant B cells develop after Vax 2 but not after Vax 1 when *H* is small because selection becomes permissive when antigen concentration is high. **(F)** *p*_1_, the fraction of positively selected GC B cells that exit GC, and *p*_2_, the fraction of such cells that become plasma cells, are varied. Left panel) If *p*_2_ increases, GC-derived B cells contribute more to the antibody titer at long times after Vax 2. This makes the antibody dynamics not consistent with clinically observed behavior where the antibody titers decay over time after Vax 2. Middle panel) If *p*_1_ increases, the ratio between GC-derived memory cells and EGC-derived memory cells changes, and the total number of memory cells increase over time. Right panel) The qualitative finding of the study is highly robust to changes in *p*_1_ and *p*_2_. Only a relatively narrow range of *p*_1_ and *p*_2_ values will be consistent with clinically observed dynamics of antibody titer and memory cell numbers, and these uncertainties will not affect the general findings of the study.

**Figure S4.**
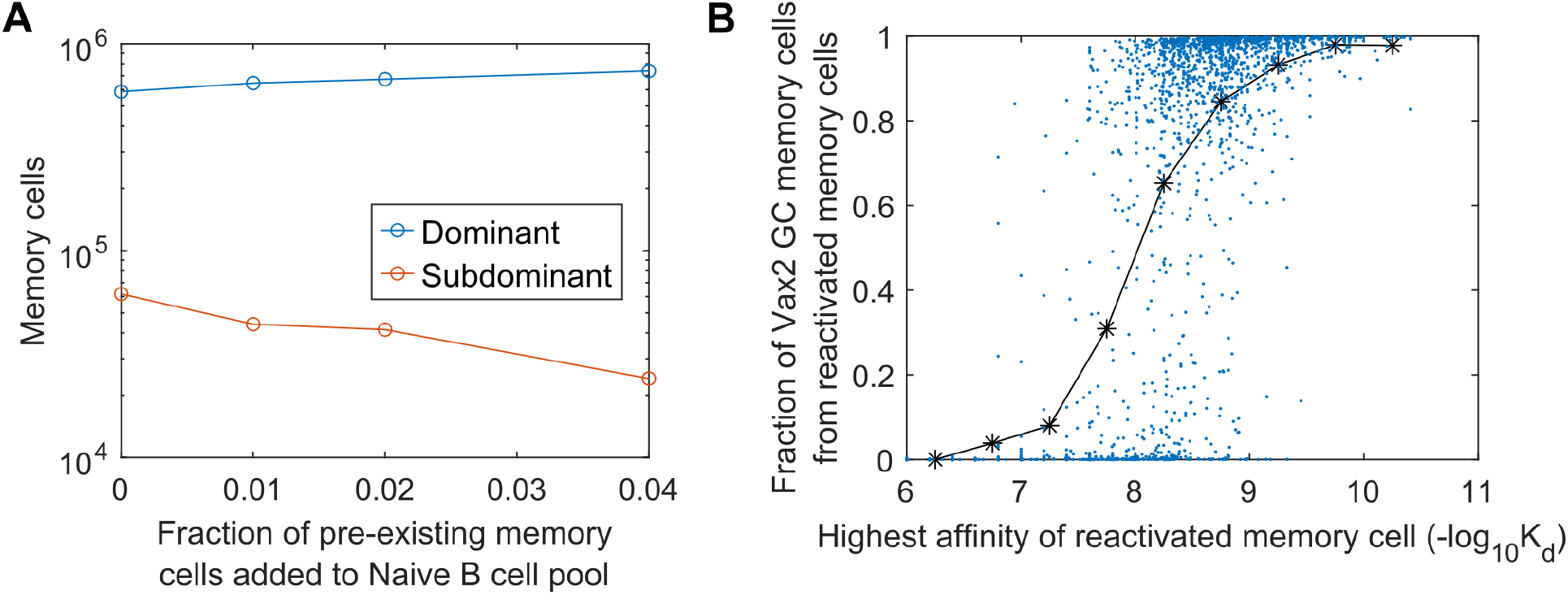
Effect of memory B cell re-entry in secondary GCs: **(A)** Number of memory B cells derived from GCs at 5 months after Vax 2 that target dominant and subdominant epitopes, when different fractions of pre-existing memory cells generated from Vax 1 GCs were allowed to re-enter Vax 2 GCs. **(B)** Fraction of memory cells derived from Vax 2 GC that are descendants of memory cells generated from Vax 1 and re-entered Vax 2 GC, as a function of the highest affinity of such re-activated memory cells. Each GC is represented by a blue dot (n=2000). The black curve shows the mean values. The fraction of pre-existing memory cells allowed to re-enter the secondary GCs is 0.04.

**Figure S5.**
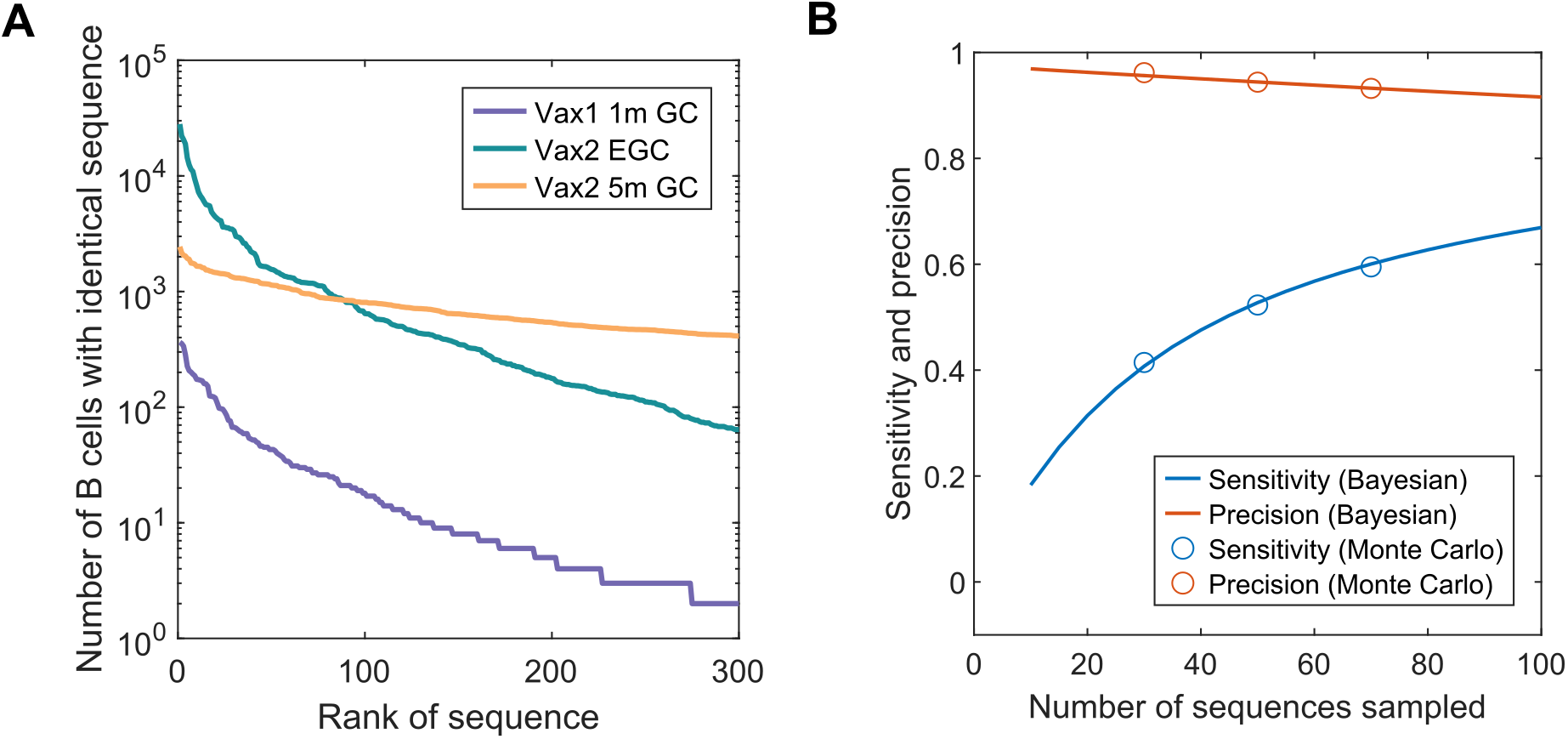
Performance of the EGC B cell labeling method: **(A)** Number of memory B cells from simulated data that have identical sequences, shown in the order of most to least expanded sequences. A few sequences dominate the Vax 2 EGC-derived memory cells. In contrast, diverse sequences of similar sizes make up the Vax 2 GC-derived memory cells. The result is from a single simulation of 200 GCs. **(B)** Sensitivity and precision of our method for finding EGC-derived B cells tested on simulated data while assuming varying numbers of sequences were sampled. The statistics calculated with Bayesian inference and with a Monte Carlo method agree well. When only a small number of sequences are sampled, the sensitivity will be low, but the precision will be high. This is likely the case for the clinical data; however, since the actual number of memory B cells in vaccinated humans will be different from the simulated data, the quantitative numbers can be different. Sensitivity is defined as (TP/TP+FN), and precision is defined as (TP/TP+FP). TP: True Positive (EGC B cell labeled as EGC), FN: False Negative (EGC B cell labeled as GC), FP: False Positive (GC B cell labeled as EGC).

**Figure S6.**
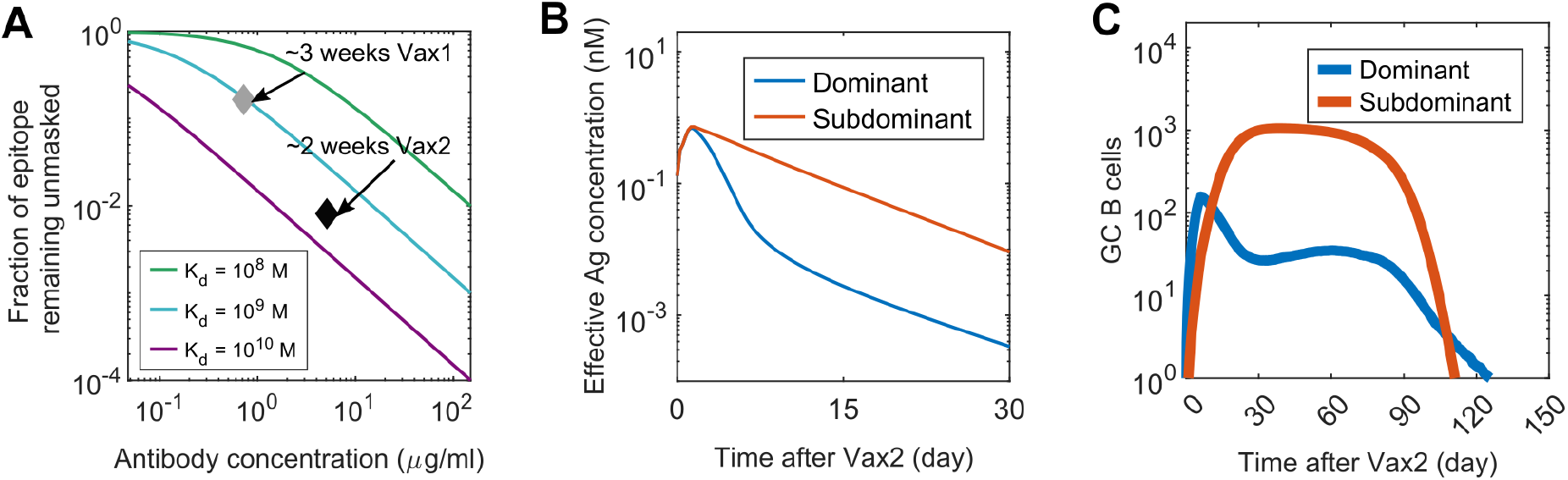
Epitope masking: **(A)** Fraction of antigen molecules that will remain unmasked at ~3 weeks after Vax 1 and ~2 weeks after Vax 2, calculated using the concentration and affinity of serum antibody from mRNA-vaccinated subjects (Demonbreun et al., 2021; Macdonald et al., 2022). **(B)** Effective antigen concentrations for B cells that target dominant and subdominant epitopes after Vax 2, when epitope masking is completely epitope-specific with no overlap. **(C)** Number of GC B cells that target dominant and subdominant epitopes after Vax 2, when epitope masking is completely epitope-specific with no overlap.

**Table S1.** Equations and parameters for antigen dynamics: Reactions that govern antigen dynamics and the differential-algebraic equations that describe the changes in concentrations of species. Initial values and parameter values are also shown. Abbreviations: soluble antigen (Ag), soluble antibody (Ig), soluble immune complex (IC), immune complex on follicular dendritic cell (IC-FDC), plasma cell (PC).

| Reaction | Description |  |
| --- | --- | --- |
| $Ag \rightarrow \emptyset$ | Decay of free soluble antigen | |
| $Ag + Ig \rightleftharpoons IC$ | Fast equilibrium between free soluble antigen and antibody | |
| $IC \rightarrow IC - FDC$ | Immune complex transport to follicular dendritic cells | |
| $PC \rightarrow PC + Ig$ | Antibody production by plasma cells | |
| $IC - FDC \rightarrow \emptyset$ | Consumption and decay of immune complexes on follicular dendritic cells | |
| $Ig \rightarrow \emptyset$ | Decay of free soluble antibody | |
| Governing Equation | Initial Condition / parameters | Note |
| $\frac{[Ag][Ig]}{[IC]} = K_d$ | $[Ag]_0 = 10 \text{ nM}$ | Values were picked within reasonable physiological ranges (Demonbreun et al., 2021; Martin et al., 2021); |
| | $[Ig]_0 = 10^{-2} \text{ nM}$ | |
| | $[IC]_0 = 0 \text{ nM}$ | Initially no IC exists |
| | $K_{d0} = 10^{-6} \text{ M}$ | Initial value for low affinity; Changes over simulation |
| $\frac{d[Ag]}{dt} = -d_{Ag}[Ag]$ | $d_{Ag} = 3 \text{ day}^{-1}$ | Picked to be fast (Aung et al.; Martin et al., 2021; Tam et al., 2016) |
| $\frac{d[IC]}{dt} = -k_{deposit}[IC]$ | $k_{deposit} = 1 \text{ hour}^{-1}$ | Picked to be fast (Aung et al.; Martin et al., 2021) |
| $\frac{d[IC - FDC]}{dt} = k_{deposit}[IC] - d_{IC}[IC - FDC]$ | $[IC - FDC]_0 = 0$ | Initially no IC-FDC exists |
| | $d_{IC} = 0.15 \text{ day}^{-1}$ | Represents the consumption and decay of antigen on FDC. Picked so that secondary GCs last ~3 months. |
| $\frac{d[Ig]}{dt} = k_{Ig}[PC] - d_{Ig}[Ig]$ | $k_{Ig} = 0.8 \times 10^{-2} \text{ nM day}^{-1} \text{ PC}^{-1}$ | Picked to match the antibody titers at the peak of Vax2 response to the values described in the literature (Goel et al., 2021; Muecksch et al., 2022). |
| | $d_{Ig} = 0.025 \text{ day}^{-1}$ | Picked to give antibody half-life of ~28 days (Goel et al., 2021) |
| $\frac{dK_d}{dt} = \frac{(K_d^{PC} - K_d)k_{Ig}[PC]}{[Ig] + [IC]}$ | $K_d^{PC}$ is the mean affinity of PCs | See star methods Eq. 1 for derivation |

**Table S2.** Simulation parameters: Description of the parameters used in the simulations. Entries highlighted with color denote the parameters whose values are varied for the robustness tests in the supplemental figures.

| Parameter | Equation | Description | Value | Note |
| --- | --- | --- | --- | --- |
| **B cells** |  |  |  |  |
| $N_{naive}$ | STAR Methods Eqs. 2-3 | Number of naïve B cells / GC | 2000 cells/GC | About $1 \times 10^{10}$ total naïve B cells (Boyd and Joshi, 2014; Rees, 2020) multiplied by the frequency of RBD-specific naïve B cells in humans is about 1 in $3 \times 10^4$ (Feldman et al., 2021) divided by 200 GCs |
| $p$ | STAR Methods Eqs. 2-3 | Fraction of germline B cells that are subdominant | 0.2 for main panels; varied between 0.02 and 0.5 for robustness test | Varied in simulation |
| $E_1^h$ | STAR Methods Eqs. 4-5 | Affinity at which there is one dominant naïve B cell available for each GC on average | 7 for main panels; varied between 7 and 8 for robustness test | Varied in simulation |
| $dE_{12}$ | STAR Methods Eq. 5 | $E_1^h - dE_{12}$ is the affinity at which there is one subdominant naïve B cell available for each GC on average | 0.4 for main panels; varied between 0 and 1 for robustness test | Varied in simulation |
| $n_{res}$ | STAR Methods Eqs. 6-7 | Length of the string representation of B cell residues | 80 | Upper range of the sum of CDR lengths in heavy and light chain (Nowak et al., 2016) |
| $\mu, \sigma, \epsilon$ | STAR Methods Eq. 8 | Parameters for the shifted log-normal distribution that represent the effects of mutations on B cell binding affinities | 3.1, 1.2, 3.08 | Fitted to empirical distribution (Zhang and Shakhnovich, 2010) |
| $\rho$ | STAR Methods Eq. 9 | Level of conservation of an epitope on WT and variant | 0.95 for subdominant, 0.4 for dominant | Picked from several values tested; no changes in qualitative findings when varied within reasonable range |
| **GC and EGC** |  |  |  |  |
| $C_0$ | STAR Methods Eq. 10 | Reference antigen concentration | $8 \times 10^{-3} nM$ for main panels; varied between $1 \times 10^{-3}$ and $6.4 \times 10^{-2} nM$ for robustness test | Varied in simulation |
| $E_0$ | STAR Methods Eq. 10 | Reference binding affinity | 6 | Corresponds to $-\log_{10} K_d = 6$, which is threshold for activation (Batista and Neuberger, 1998) |
| $K$ | STAR Methods Eq. 10 | Stringency of selection of naïve and GC B cells by helper T cells based on the amounts of antigen internalized | 0.5 for main panels; varied between 0.3 and 1 for robustness test | Varied in simulation |
| $N_{max}$ | STAR Methods Eq. 12 | Approximately the maximum number of naïve B cells that can enter GC per day | $10 \text{ day}^{-1}$ | Picked to match experimental observation in mice (Tas et al., 2016) |
| $H$ | STAR Methods Eq. 14 | Parameter that controls the saturation point when alternative definition for antigen internalization is used | 20, 100, 500 | Varied in simulation |
| $\beta_{max}$ | STAR Methods Eq. 15 | Maximum rate of positive selection for GC and EGC B cells | $2.5 \text{ day}^{-1}$ | Maximum proliferation is about ~4 times / day (Victora et al., 2010) |
| $N_{T0}$ | STAR Methods Eq. 17 | Maximum number of helper T cells | 1200 | Picked to give peak GC size of ~1000 B cells |
| $t_0$ | STAR Methods Eq. 17 | Time at which the number of helper T cells reaches maximum | 14 day | Matches the dynamics in mRNA-vaccinated humans (Goel et al., 2021) |
| $d_T$ | STAR Methods Eq. 17 | Rate of decay for helper T cells | $0.015 \text{ day}^{-1}$ | Matches the dynamics in mRNA-vaccinated humans (Goel et al., 2021) |
| $\alpha$ | STAR Methods Eq. 18 | Death rate of GC B cells | $0.5 \text{ day}^{-1}$ | Picked to allow B cells to survive ~2 days before death if they don't get selected |
| **Memory and Plasma Cell Dynamics** |  |  |  |  |
| $p_1$ | - | Probability that a positively selected GC B cell exits by differentiation | 0.05 for main panels; varied between 0.03 and 0.1 for robustness test | Varied in simulation |
| $p_2$ | - | Probability that a differentiating GC B cell becomes a plasma cell | 0.1 for main panels; varied between 0.1 and 0.5 for robustness test | Varied in simulation |
| $p_2^{EGC}$ | - | Probability that a proliferating memory cell in EGC differentiates into a plasma cell | 0.6 | Based on observation in mice (Moran et al., 2018) |
| $d_{PC}$ | STAR Methods Eq. 19 | Death rate of plasma cells | $0.17 \text{ day}^{-1}$ | Short-lived plasma cells have half life of ~4 days (Khodadadi et al., 2019) |

**Table S3.**
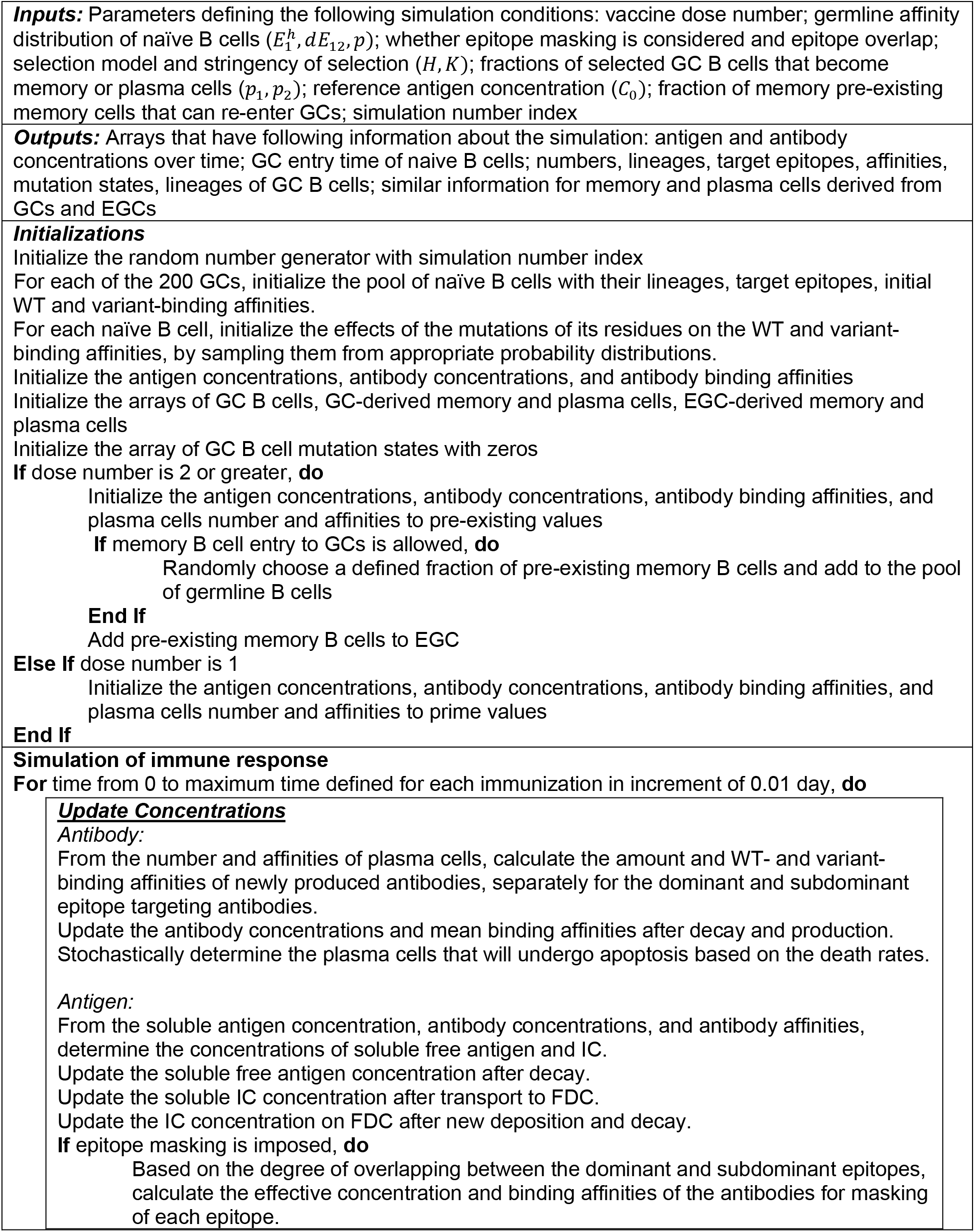
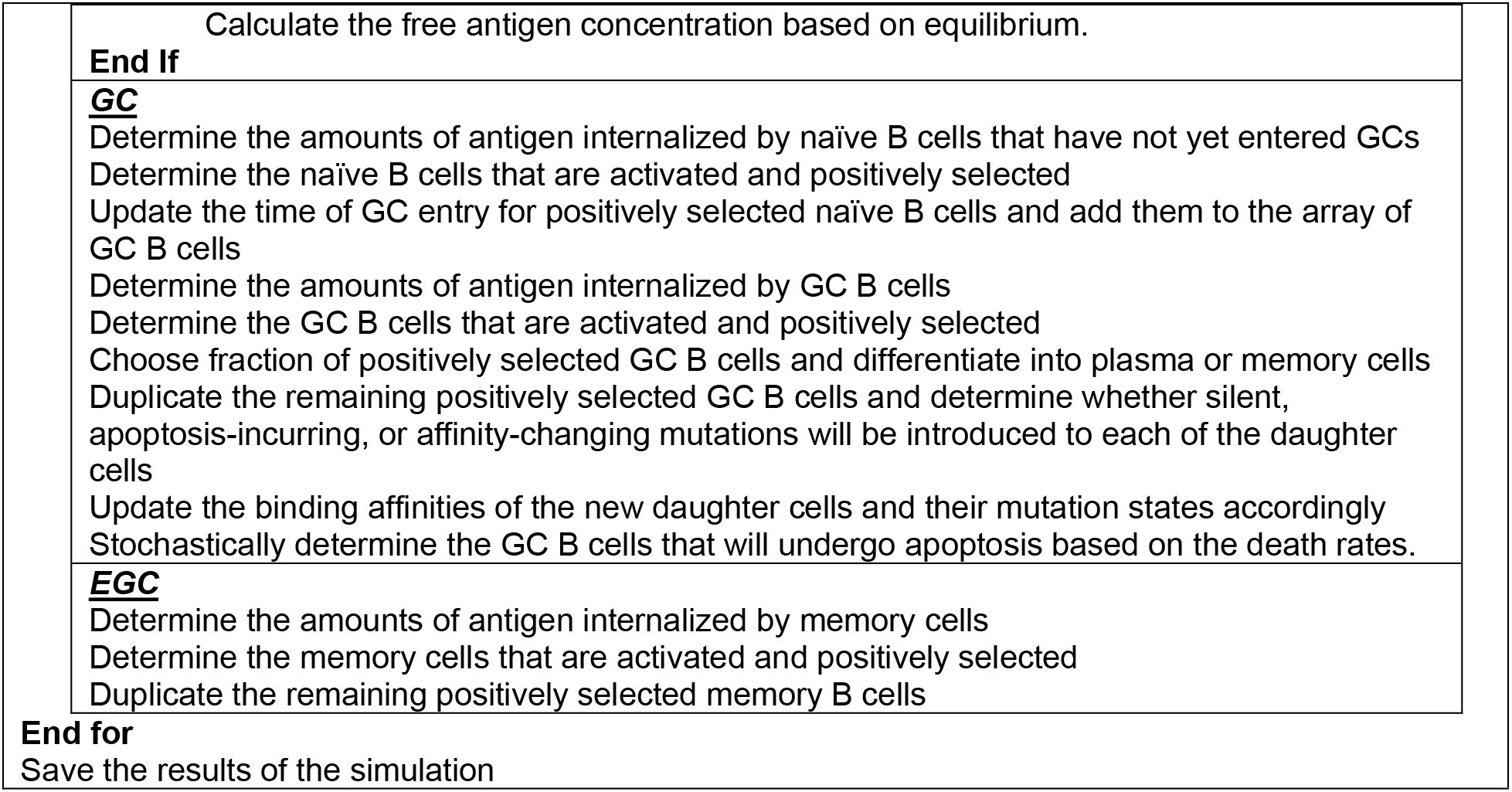
Summary of the simulation algorithm: Pseudocode that summarizes the simulation algorithm implemented in MATLAB. It describes a single run of simulation that models 200 GCs and an EGC.

|  |
| --- |
| **Inputs:** Parameters defining the following simulation conditions: vaccine dose number; germline affinity distribution of naïve B cells ($E_1^h, dE_{12}, p$); whether epitope masking is considered and epitope overlap; selection model and stringency of selection ($H, K$); fractions of selected GC B cells that become memory or plasma cells ($p_1, p_2$); reference antigen concentration ($C_0$); fraction of memory pre-existing memory cells that can re-enter GCs; simulation number index |
| **Outputs:** Arrays that have following information about the simulation: antigen and antibody concentrations over time; GC entry time of naïve B cells; numbers, lineages, target epitopes, affinities, mutation states, lineages of GC B cells; similar information for memory and plasma cells derived from GCs and EGCs |
| **Initializations**<br>Initialize the random number generator with simulation number index<br>For each of the 200 GCs, initialize the pool of naïve B cells with their lineages, target epitopes, initial WT and variant-binding affinities.<br>For each naïve B cell, initialize the effects of the mutations of its residues on the WT and variant-binding affinities, by sampling them from appropriate probability distributions.<br>Initialize the antigen concentrations, antibody concentrations, and antibody binding affinities<br>Initialize the arrays of GC B cells, GC-derived memory and plasma cells, EGC-derived memory and plasma cells<br>Initialize the array of GC B cell mutation states with zeros<br>**If** dose number is 2 or greater, **do**<br>Initialize the antigen concentrations, antibody concentrations, antibody binding affinities, and plasma cells number and affinities to pre-existing values<br>**If** memory B cell entry to GCs is allowed, **do**<br>Randomly choose a defined fraction of pre-existing memory B cells and add to the pool of germline B cells<br>**End If**<br>Add pre-existing memory B cells to EGC<br>**Else If** dose number is 1<br>Initialize the antigen concentrations, antibody concentrations, antibody binding affinities, and plasma cells number and affinities to prime values<br>**End If** |
| **Simulation of immune response**<br>**For** time from 0 to maximum time defined for each immunization in increment of 0.01 day, **do**<br>**<u>Update Concentrations</u>**<br>**Antibody:**<br>From the number and affinities of plasma cells, calculate the amount and WT- and variant-binding affinities of newly produced antibodies, separately for the dominant and subdominant epitope targeting antibodies.<br>Update the antibody concentrations and mean binding affinities after decay and production.<br>Stochastically determine the plasma cells that will undergo apoptosis based on the death rates.<br>**Antigen:**<br>From the soluble antigen concentration, antibody concentrations, and antibody affinities, determine the concentrations of soluble free antigen and IC.<br>Update the soluble free antigen concentration after decay.<br>Update the soluble IC concentration after transport to FDC.<br>Update the IC concentration on FDC after new deposition and decay.<br>**If** epitope masking is imposed, **do**<br>Based on the degree of overlapping between the dominant and subdominant epitopes, calculate the effective concentration and binding affinities of the antibodies for masking of each epitope. |
| Calculate the free antigen concentration based on equilibrium. |
| **End If** |
| **<u>GC</u>** |
| Determine the amounts of antigen internalized by naïve B cells that have not yet entered GCs<br>Determine the naïve B cells that are activated and positively selected<br>Update the time of GC entry for positively selected naïve B cells and add them to the array of GC B cells<br>Determine the amounts of antigen internalized by GC B cells<br>Determine the GC B cells that are activated and positively selected<br>Choose fraction of positively selected GC B cells and differentiate into plasma or memory cells<br>Duplicate the remaining positively selected GC B cells and determine whether silent, apoptosis-incurring, or affinity-changing mutations will be introduced to each of the daughter cells<br>Update the binding affinities of the new daughter cells and their mutation states accordingly<br>Stochastically determine the GC B cells that will undergo apoptosis based on the death rates. |
| **<u>EGC</u>** |
| Determine the amounts of antigen internalized by memory cells<br>Determine the memory cells that are activated and positively selected<br>Duplicate the remaining positively selected memory B cells |
| **End for** |
| Save the results of the simulation |

**Table S4.** Neutralization activities of the recombinant antibodies derived from the memory B cells identified as EGC-derived: IC50s against Omicron were measured using Omicron HT1080/Ace2 cl14 cells. IC50s against WT were measured using WT(R683G) HT1080/Ace2 cl14 cells unless otherwise noted. The IC50s against Omicron and WT were measured newly for this study except for C3136, C3050, and C2593, whose IC50 values are taken from previously reported values (Muecksch et al., 2022). More information about the antibodies including their sequences, germline gene usage, and somatic mutations can be found in the supplemental tables of Muecksch et al. based on their IDs (Muecksch et al., 2022).

| Antibody ID | Found In | IC50 Omicron | IC50 WT | Counted As EGC Clone in |
| --- | --- | --- | --- | --- |
| **C2107** | Vax1/Vax2 1.3m | 1000.0 | 1000.0 | Vax2 |
| **C2173** | Vax2 1.3m | 1000.0 | 20.4 | Vax2 |
| **C4002** | Vax2 1.3m | 1000.0 | 1000.0 | Vax2 |
| **C2175** | Vax2 1.3m | 918.9 | 11.6 | Vax2 |
| **C2174** | Vax2 1.3m | 920.4 | 347.2 | Vax2 |
| **C2478** | Vax2 1.3m/5m | 948.5 | 1000.0 | Vax2 |
| **C3137** | Vax2 5m | 694.8 | 1000.0 | Vax2 |
| **C3136\*** | Vax2 5m | 896.7 | 28.5** | Vax2 |
| **C3050\*** | Vax2 5m/Vax3 | 931.3 | 1000** | Vax2 and Vax3*** |
| **C3138** | Vax2 5m/Vax3 | 522.5 | 8.0 | Vax3 |
| **C2593\*** | Vax3 | 0.9 | 3.9** | Vax3 |
| **C4001** | Vax3 | 7.4 | 0.9 | Vax3 |
| **C4003** | Vax3 | 300.6 | 21.6 | Vax3 |
| **C3109** | Vax3 | 271.0 | 77.8 | Vax3 |
| **C3112** | Vax3 | 44.7 | 27.3 | Vax3 |
| **C3117** | Vax3 | 36.4 | 9.8 | Vax3 |

\*The neutralizing activities of C3136, C3050, C2593 are taken from Muecksch et al.
\*\*IC50 values for these antibodies were measured against the WT 293TAce2 cells, as reported in Muecksch et al. (2022)
\*\*\*Multiple identical sequences were found at each timepoint

## STAR METHODS

### RESOURCE AVAILABILITY

- Lead Contact
- Materials availability
- Data and code availability

### METHODS DETAILS

- Simulation Details for Antigen Transport and Presentation
- Simulation Details for B cells and 2-epitope model
- Alternative Model for Antigen Capture
- Simulation Details for GCs
- Simulation Details for EGC
- Clinical Sample Collection and Analysis Methods
- Sensitivity and Precision of the Inference of EGC-derived Memory Cells
- Epitope Masking

### RESOURCE AVAILABILITY

#### Lead contact

Further information and requests for resources and reagents should be directed to and will be fulfilled by the lead contact, Arup K. Chakraborty.

#### Materials availability

This study did not generate new unique reagents.

#### Data and code availability

- Simulation data have been deposited at github.com/leerang77/Booster_Shot_Variant and are publicly available.
- All original code has been deposited at github.com/leerang77/ Booster_Shot_Variant and is publicly available.
- Any additional information required to reanalyze the data reported in this paper is available from the lead contact upon request.

### METHOD DETAILS

#### Simulation Details for Antigen Dynamics

Table S1 describes the reactions that govern antigen dynamics and the differential-algebraic equations derived from the reactions that are solved in the simulations. The values of initial conditions and parameters are also shown, with notes on how they were selected. The following species are involved in the dynamics: soluble antigen (Ag), soluble antibody (Ig), soluble immune complex (IC), immune complex on follicular dendritic cell (IC-FDC), and plasma cell (PC).

The simulation progresses in time steps of 0.01 day, and the concentrations are updated at each step. Since the on-rate for antigen and antibody binding is very fast (order of *K*_*a*_ = 10^11^ *M*^−1^*day*^−1^) (Batista and Neuberger, 1998), we assume that fast equilibrium is maintained between Ag, Ab, and IC. Thus, the equilibrium concentrations [Ag], [Ig], and [IC] can be calculated. Then, the concentrations of all species except for the PCs are updated to account for Ag decay, IC deposition on FDC, Ig production by plasma cells, IC-FDC consumption, and Ig decay, based on the differential equations described in Table S1. The PC concentration is updated based on their stochastic production and apoptosis from B cell dynamics involving GCs and EGCs. Each simulation models 200 GCs and 1 EGC simultaneously, and all the PCs derived from them contribute to the Ig kinetics. After all of the concentrations are updated at each step, the mean antibody association constant *K*_*a*_ for the WT and the variant are updated. The governing equation is derived using the product rule as follows:

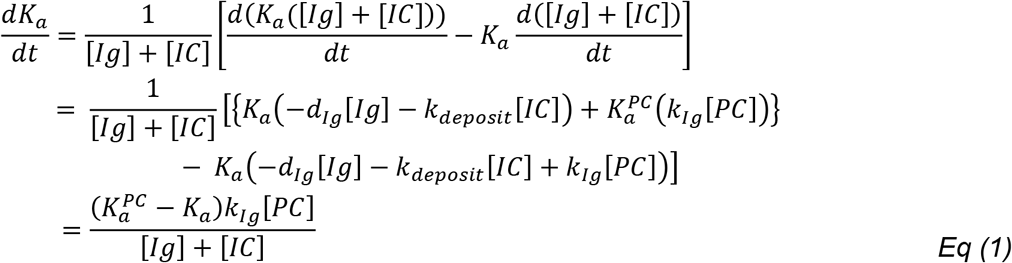

*K*_*a*_ and 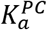 are the mean association constants of the existing antibodies and PCs, respectively, and *K*_*Ig*_ is the rate of antibody production per plasma cell. The other parameters are described in Table S1. The derivation makes use of the fact that the change in total antibody titer, *K*_*d*_ ([*Ig*] + [*IC*]), can be obtained from the consumption and production of the antibody species.

For Vax 1, the initial concentrations for IC, IC-FDC, and PC are set to zeros and the initial concentration for Ag is set to 10 *nM* to represent a bolus injection of antigen. There will be only a small number of weakly-binding antibodies to the new immunogen, so [*Ig*] and *K*_*d*_ are initially set to small values. These values and other parameters in Table S1 are picked from reasonable physiological ranges based on the literature (Demonbreun et al., 2021; Macdonald et al., 2022; Martin et al., 2021; Tam et al., 2016). While there are uncertainties about the true underlying biological values, the physical significances of these initial values and parameters are in determining the level of antigen availability in the lymph node. In our model, the antigen availability depends on the reference antigen concentration *C*_0_ because antigen capture by B cells depends on the normalized antigen availability 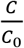, where *C* is the amount of antigen in the *c*0 lymph node. Thus, by changing *C*_0_, we can study the effect of changing antigen availability in the system. As mentioned in the main figures and shown in Fig S4A, we tested the robustness of the results on varying *C*_0_.

For Vax 2, Vax 3, and Vax 4, the initial concentrations of the species are set to the values determined by response to the previous vaccination.

#### Simulation Details for B cells and 2-Epitope Model

As described in the main text, the dynamics of B cells are simulated with an agent-based model. Each B cell is an agent that has the following properties: type, lineage, target epitope, mutational state, and binding affinities. At each time point, the B cells stochastically undergo different actions based on their properties and the conditions of the simulation. The details of the model are described below, and the simulation algorithm is summarized in Table S3. Table S2 summarizes the parameters used in the simulations. It shows which equations the parameters appear in, their descriptions, values, and notes about how those values were selected.

Each simulation models 200 GCs simultaneously. Each GC is associated with a pool of naïve B cells that have not yet entered the GC. The number of total naïve B cells in humans is estimated to be about 1 × 10^10^ (Boyd and Joshi, 2014; Rees, 2020), and the frequency of SARS-CoV-2 RBD-specific naïve B cells is about 1 in 3 × 10^4^ (Feldman et al., 2021). Thus, we assume that the number of naïve B cells for each GC is *N*_*naive*_ = (1 × 10^10^)/(3 × 10^4^) / 200 ≅ 2000 cells. These naïve B cells have germline-endowed WT-binding affinities, whose possible values are *E*_*K*_ = 6 + 0.2*K* (*K* = 0…10). These affinities correspond to − log_10_ *K*_*d*_. The distribution of the naïve B cells over the possible values is determined by three parameters: 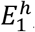, *dE*_12_, *p*. Higher-affinity B cells should be rarer, so the frequency of B cells is determined analogously to a truncated geometric distribution (see Figure S1A for the schematics). The frequency of naïve B cells that target the dominant and subdominant epitopes are as follows:

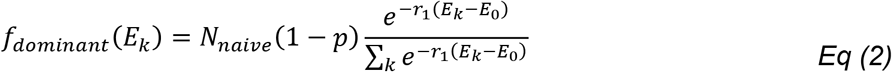

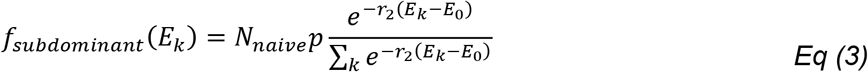

*p* is the fraction of naïve B cells that target the subdominant epitope, and *r*_1_, *r*_2_ in the exponents are specified by the parameters 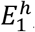 and *dE*_12_ from the following relationships.

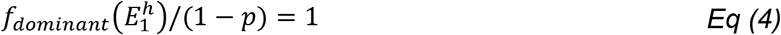

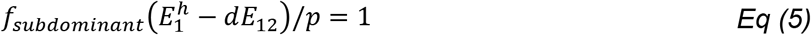

That is, 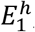 and 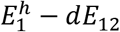 are the affinities at which the frequency of naïve B cells that target the dominant and subdominant epitopes respectively would be 1 cell per GC, before adjusting for the total frequency (Fig S1A). For each GC, the exact number of naïve B cells that have germline affinity equal to *E*_*K*_ is determined by stochastically rounding up or rounding down *f*_*dominant*_(*E*_*K*_) and *f*_*subdominant*_(*E*_*K*_) to the nearest integer, using the fractional part as the probability of rounding up. Very high-affinity naïve B cells have precursor frequencies of less than 1 per GC (Figure S1A), so they will exist only for some of the GCs.

Each naïve B cell also has a germline-endowed binding affinity against the variant strain. Immunization with the WT strain will recruit naïve B cells with high WT-binding affinities; even the naïve B cells with the lowest WT-binding affinity in the pool (*E*_0_ = 6) still represents the top 1 in ~3 × 10^4^ of all naïve B cells in the human repertoire. The binding affinity of these naïve B cells against the variant will likely be lower. Thus, we assume that all naïve B cells have germline binding affinity of − log_10_ *K*_*d*_ = 6 against the variant, equal to the lowest value of binding affinity against the WT, and that required for GC entry (Batista and Neuberger, 1998).

During affinity maturation, the affinities of B cells change as they accumulate mutations. To account for mutations, each naïve B cell is represented as a string of 0’s with length *N*_*vvdddd*_, and an affinity-affecting mutation to a GC B cell changes the value of one randomly selected residue from 0 to 1 or from 1 to 0. Each residue that has a value of 1 changes the binding affinity towards the WT and the variant by pre-determined amounts. These amounts, which are analogous to the fitness landscape of the B cell, are drawn from a correlated probability distribution. Fig. S1B schematically shows how the affinities are determined for GC B cell, *i*. The binding affinities against the WT and the variant are determined as

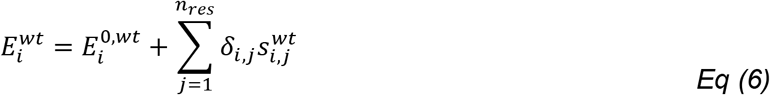

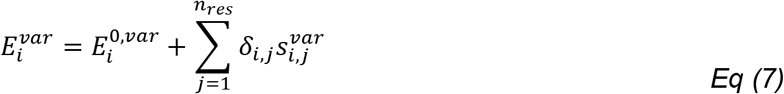

where 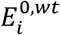 and 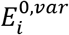 are the germline affinities towards the WT and the variant, respectively; *δ*_*i*,*j*_ ∈ {0,1} is the mutational state of residue *j*; and 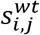 and 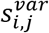 are the effects of the mutation at residue *j* on the binding affinities against the WT and the variant, respectively. 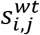 and 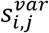 are sampled from the following shifted log-normal distribution, independently for each residue *j*, at the initiation of the simulation.

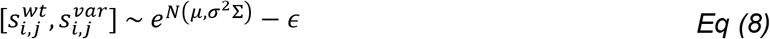

The parameters *μ*, *σ*, *ϵ* are chosen to fit experimentally determined distribution, where ~5 % of affinity-affecting mutations are beneficial while most of the mutations are strongly deleterious (Figure S1C) (Kumar and Gromiha, 2006; Zhang and Shakhnovich, 2010). The covariance has the form

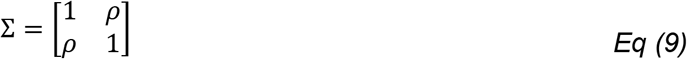

where *ρ* represents the level of conservation of an epitope between the WT and variant. As *ρ* increases, mutations that are beneficial for binding both strains become more common (Figure S1D). We choose *ρ* = 0.95 for the subdominant epitope and *ρ* = 0.4 for the dominant epitope. For B cells that target the subdominant and dominant epitope, respectively 72% and 19% of mutations that are beneficial for binding the WT are also beneficial for binding the variant, and vice versa. Since B cells are selected in GCs based on their WT-binding affinities, an increase in variant-binding affinities mainly occurs through the accumulation of mutations that increase affinities against both strains. Hence, B cells that target the subdominant epitope are more likely to develop high cross-reactivity for the variant than those that target the dominant epitope.

#### Simulation Details for Germinal center entry of naïve B cells

At each time step, the amount of antigen captured by naive B cells is determined based on their WT-binding affinities and the effective antigen concentration in the lymph node, *C*. For B cell *i*, this amount, *A*_*i*_, is determined as follows:

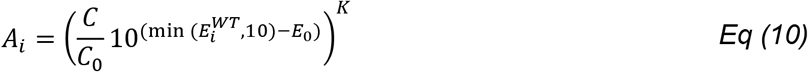

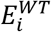 is the WT-binding affinity of B cell *i*. The amount of antigen captured increases with 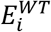, but saturates at affinities higher than 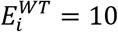 because of the affinity ceiling (Foote and Eisen, 1995). A similar model of antigen capture has been used in several previous studies (Amitai et al., 2017, 2020; Molari et al., 2020; Wang et al., 2015). B cells can see both the soluble antigen and the antigen presented on FDCs, but the latter is known to be about 2 orders of magnitude more potent at activating B cells due to multivalent presentation (Kim et al., 2006). Therefore, the effective antigen concentration *C* is calculated as *C* = 0.01([*Ag*] + [*IC*]) + [*IC* − *FDC*]. The parameter *K* determines how much a given difference in concentration or affinity changes the amount of antigen internalized by a B cell. If *K* is large, then even a small difference in concentration or affinity results in large difference in the amount of antigen internalized, which in turn affects the probability of activation and positive selection by T helper cells. Thus, *K* represents the stringency of selection. We studied varying *K* to test the robustness of the results, since stringency of selection is known to affect the diversity of B cells that develop in GCs (Victora and Wilson, 2015) (Fig S3D).

Naïve B cells that capture enough antigen can be activated (Batista and Neuberger, 1998). In our simulation, whether B cell *i* is activated at each time step is determined probabilistically as follows:

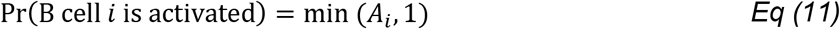

The entry of activated naïve B cells to GCs is limited by competition for positive selection by helper T cells, and B cells that have internalized greater amounts of antigen have better chances of successfully entering GCs (Lee et al., 2021; Schwickert et al., 2011). Thus, the rate of entry for an activated B cell *i*, *λ_i_*, and the probability that it enters GC during a time step are given as follows:

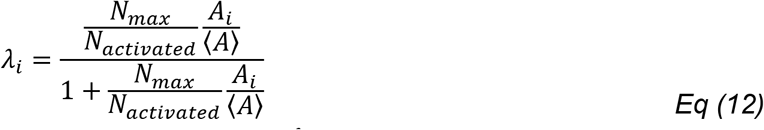

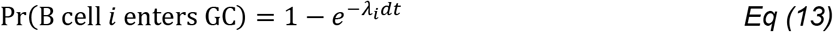

*N*_*activated*_ is the total number of activated B cells, 〈*A*〉 is the average amount of antigen captured by all activated B cells, and *N*_*max*_ is the capacity for entry that represents the limited amount of T cell help. *N*_*max*_ is selected so that about ten distinct naïve B cells will enter the GC per day, consistent with the literature (Tas et al., 2016). The assumption that *N*_*max*_ is fixed is conservative because higher antigen availability is known to increase naïve B cell recruitment to GCs (Angeletti et al., 2019), which would only further strengthen our finding that secondary GCs produce more diversity. When a naïve B cell enters GC, it simultaneously proliferates twice, so that a total of 4 identical B cells are added to the GC.

#### Alternative Model for Antigen Capture

According to Eq. 10, the amount of antigen captured by B cells continues to increase with antigen concentration and B cell affinity. However, it is possible that the amount of antigen captured plateaus when antigen concentration and B cell affinity are very high (Fleire et al., 2006). Therefore, we studied how using an alternative model where antigen capture saturates at high affinities and antigen concentrations affects our findings. Under this model, the amount of antigen captured is determined as:

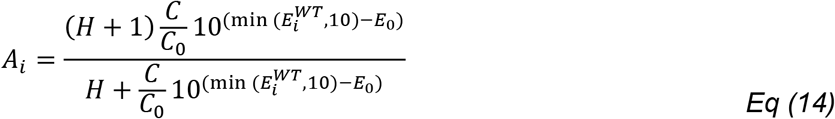

When *H* → ∞, this formulation becomes equivalent to Eq. 10 with K=1. For a finite value of *H*, *A*_*i*_ saturates to *H* + 1 when 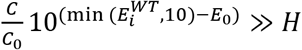. When *H* is smaller and antigen availability is higher, the affinity at which saturation will occur will be lower, making the selection of B cells permissive. We studied the effect of varying *H* on our findings (Fig. S4E).

#### Simulation Details for GCs

Each simulation models 200 GCs simultaneously. Plasma cells generated from all GCs collectively determine antibody production, which affects antigen transport and epitope masking, and memory B cells generated from all GCs seed the EGC upon subsequent vaccination. The birth, death, mutation, and differentiation of GC B cells occur stochastically at each time step. The model does not have a spatial resolution of the GC light zone and dark zone recycling, but such a model has been shown to recapitulate qualitative GC dynamics well (Amitai et al., 2017, 2020).

GC B cells capture antigen and become stochastically activated in the same way as the naïve B cells. Activated GC B cells compete for positive selection signals from helper T cells. The rate of positive selection for a GC B cell *i*, *β*_*i*_, and the probability that it gets positively selected during a time step are given as:

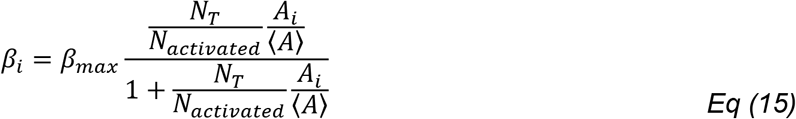

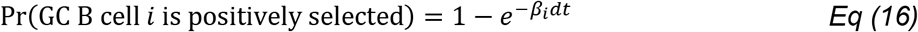

where *β*_*max*_ is the maximum rate of positive selection, *N*_*activated*_ is the number of activated GC B cells, and *N*_*T*_ is the number of helper T cells. Thus, 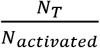 represents the physical availability of helper T cells to GC B cells, and 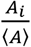 represents the competitive advantage of B cell *i* compared to other activated GC B cells.

Clinical data from SARS-CoV-2 vaccinated subjects showed that the number of CD4^+^ T cells peaked about 2 weeks after vaccination and decayed with a half-life of ~47 days (Goel et al., 2021). For simplicity, we model *N*_*T*_ as simple linear growth up to *t*_0_ = 14 *days*, followed by first-order decay afterwards with rate *d*_*T*_ as follows:

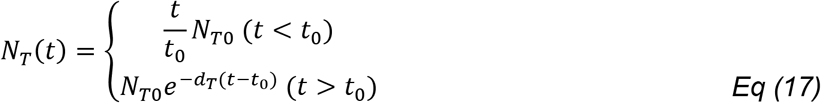

*N*_*T*0_ is the peak level of non-dimensionalized T cell availability, and is chosen to give a mean peak GC size of ~1000 cells/GC.

A positively selected B cell exits a GC with a probability *p*_1_, and then differentiates into a PC with a probability *p*_2_ or into a memory cell with a probability 1 − *p*_2_. The remaining selected B cells proliferate once and one of the daughter cells mutates, as described in the main text.

At the end of the time step, all GC B cells are subject to stochastic apoptosis with a rate *α*. The probability of apoptosis is given as:

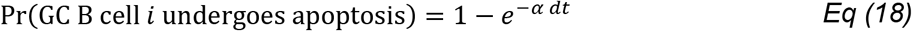

Similarly, plasma cells from both GCs and EGCs also undergo stochastic apoptosis at a rate *d*_*PC*_, so that the probability of apoptosis is given as:

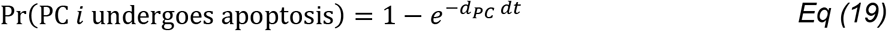

#### Clinical Sample Collection and Analysis Methods

Data used in Figure 4 are derived from B cell sequences reported in Supplemental Table 2 of Muecksch et al., which contains sequences of B cells isolated from SARS-CoV-2 mRNA-vaccinated subjects (Muecksch et al., 2022). Phylogenetic trees were generated from these B cell clonal families using MATLAB’s seqlinkage function. EGC-derived B cells were identified by applying the classification method described in the main text and in the next section. Then, using the monoclonal antibodies that correspond to these B cells based on the protein sequences (reported in Supplemental Table 3 of Muecksch et al.), the WT and Omicron-neutralizing activity (IC50) of these sequences were measured, except for three antibodies for which both values were already reported in the Supplemental Table 4 and 5 of Muecksch et al. We additionally measured the neutralization activity of 26 randomly-selected singlets that were found 5 months after Vax 2, to compare with the EGC-derived antibodies. Table S4 describes the neutralization activities of the EGC-derived antibodies used in this study.

The statistical analyses to compare the neutralization activity of EGC- and GC-derived antibodies were performed based on the logarithm of IC_50_ data. We used the two-sample t-test to calculate the statistical significance (p-value) of the difference in the mean values between the two groups. The degrees of freedom were conservatively estimated using the smaller sample size of the two samples, so that it was given as one less than the number of sequences in the smaller group. The analysis was performed to compare Vax 2 EGC-derived cells with Vax 2 GC-derived cells, and to compare Vax 2 EGC-derived cells with Vax 3 EGC-derived cells.

#### Sensitivity and Precision of the Inference of EGC-derived Memory Cells

A B cell was identified as EGC-derived if it satisfied at least one of the two conditions below.

1. Criteria 1: At least one other identical sequence was sampled at the same time
2. Criteria 2: At least one identical sequence was sampled at an earlier time

Assume that after secondary immunization, the sets of unique memory B cell sequences derived from GC and EGC are 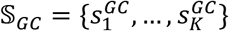 and 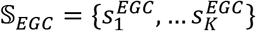}, respectively. Without the loss of generality, let the number of GC-derived memory B cells that have sequences 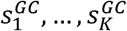 to be *m*_1_ > … > *m*_*K*_ for GC-derived cells. Similarly, let the number of EGC-derived memory B cells that have sequences 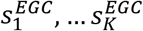 to be *n*_1_ > … > *n*_*K*_ for EGC-derived cells. *K* is a sufficiently large number. If the actual number of GC-derived unique sequences is smaller than *K*, then *n*_*i*_ will be zero for some large values of *i*. The same is true for EGC-derived sequences.

The sequences 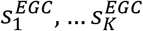 must be identical to the sequences derived from the GC of the primary immunization. Let the numbers of B cells from the prime GC that correspond to these sequences be *l*_1_, …, *l*_*K*_.

Suppose that total of *S* sequences are sampled each after the secondary immunization and the primary immunization. Let these sequences be 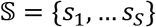 and 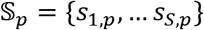, respectively. Based on the two criteria, a B cell *i* sampled after secondary immunization is labeled as EGC-derived if and only if

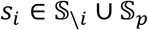

where 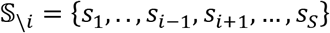 is defined as the set of sequences in 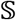 excluding *s*_*i*_.

The sensitivity, or true positive rate, of the classification is defined as the following expected value:

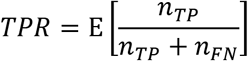

where *n*_*TP*_ and *n*_*FN*_ are the number of true positives and false negative in the labeled samples. A true positive sample is an EGC-derived sequence labeled as EGC-derived, and false positive is an EGC-derived sequence labeled as GC-derived.

An equivalent definition for sensitivity is the probability that an EGC-derived sequence will be labeled correctly as EGC-derived. That is,

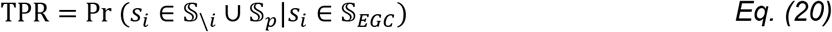

Let 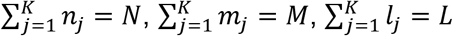. Then, the sensitivity can be calculated as

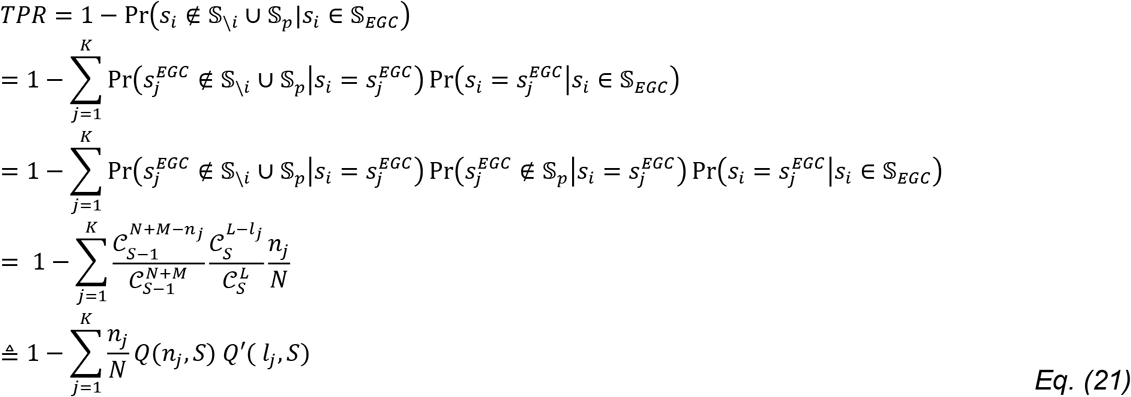

where 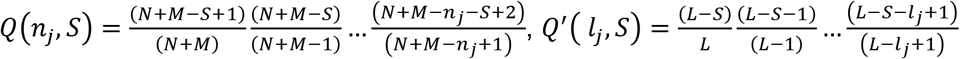

*Q*(*n*_*j*_, *S*) decreases with *n*_*j*_ and *S*. Thus, the sensitivity will be high if most B cells belong to largely expanded sequences, and if the sampling number is large. *Q*′(*l*_*j*_, *S*) decreases with *l*_*j*_ and *S*. Thus, the sensitivity will be high if for the values of *j* such that *n*_*j*_ is large, *l*_*j*_ is also large.

The precision, or positive predictive value, of the classification is defined as

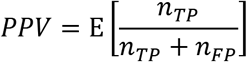

where *n*_*FN*_ is the number of false positives, or GC-derived B cells labeled as EGC-derived. An equivalent definition for precision is the probability that an EGC-labeled B cell is a true EGC-derived B cell.

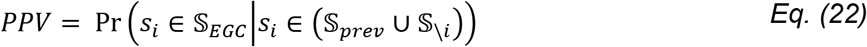

Using Bayes’ rule,

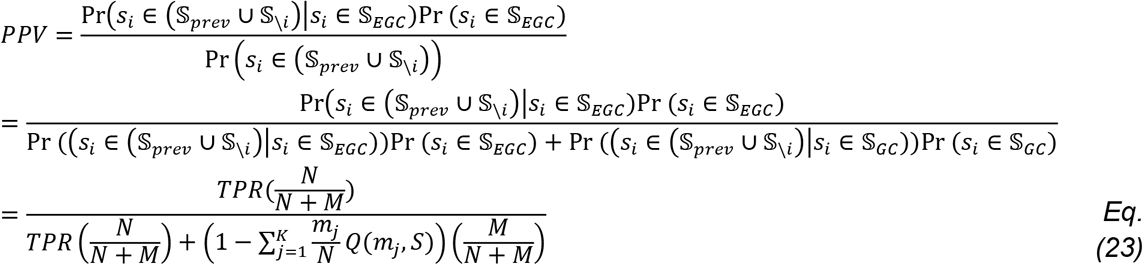

Assuming that *N* and *M* are similar, high precision is reached if the values of *Q*(*m*_*j*_, *S*) are large for the GC-derived B cells. Since *Q*(*m*_*j*_, *S*) increases with decreasing *m*_*j*_, precision is high if many GC-derived sequences have similar sizes.

We applied this analysis to the data from simulations to find the sensitivity and precision of the method. We also tested the analysis against Monte-Carlo sampling of sequences from the simulations. For this, we sampled equal numbers of memory B cells from 1 month after Vax 1 and 5 months after Vax 2. Then we applied the labeling method and calculated the number of true positives, false negatives, and false positives. This was repeated 1000 times to calculate the mean sensitivity and precision.

#### Epitope Masking

When epitope masking is considered in the simulations, B cells can only see free antigen. The total amount of antigen in a lymph node is [*Ag*] + [*IC*] + [*IC* − *FDC*] = [*Ag*]_*tot*_. Let us use subscripts 1 and 2 to denote antibodies that target the dominant and subdominant epitopes, respectively, and let *Q* be the epitope overlap. The effective concentration and mean binding affinity of the antibodies that cover the dominant epitope are [*Ig*_1_]_*eff*_ = [*Ig*_1_] + *Q*[*Ig*_2_] and 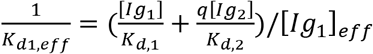, respectively. Using these values, the free antigen concentration for the dominant epitope, [*Ag*]_*tot*,1_, can be calculated from the following equilibrium.

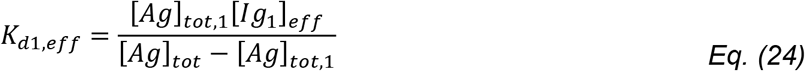

Here, we used the fact that typically [*Ig*_1_]_*eff*_ ≫ [*Ag*]_*tot*_ to approximate the free antibody concentration. Finally, to calculate the effective free antigen concentration for the dominant epitope, *C*_*eff*,1_, we must adjust for the fractions of the free antigen that are soluble or on FDC as follows:

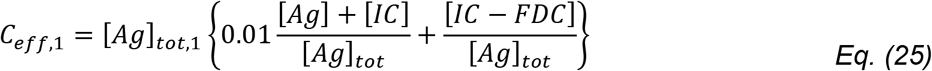

The effective free antigen concentration for the subdominant epitope can be calculated similarly. Note that although ICs are tethered to FDC, we treat them as free antigen unless it is additionally covered by serum antibody, similar to the computational model from a previous study (Zhang et al., 2013). In the experimental part of this study, mice were immunized with 4-hydroxy-nitrophenyl coupled to chicken gamma globulin (NP-CGG) along with NP-specific antibodies so that the ICs were deposited on FDCs. These ICs on FDCs elicited NP-specific serum response, suggesting that the NP epitope was not blocked by the tethering of IC to FDC.

